# Molecular clockwork hypothesis for the KaiABC circadian oscillations

**DOI:** 10.64898/2026.05.07.723666

**Authors:** Masaki Sasai, Shin Fujishiro

## Abstract

When three cyanobacterial proteins—KaiA, KaiB, and KaiC—are incubated with ATP in vitro, the phosphorylation level of KaiC exhibits stable circadian oscillations. Biochemical and structural analyses have shown that KaiC’s ATPase activity is crucial for these oscillations, leading to the hypothesis that ATP-consuming dynamics function as a molecular clock, determining the oscillation period of individual molecules. Moreover, these molecular clocks synchronize collectively, resulting in oscillations at the ensemble level. In this study, we develop a theoretical model to test this molecular clockwork hypothesis. Our model clarifies the relationship between the oscillation period and ATPase activity, explaining the significant changes in the period induced by amino acid substitutions near the CI-CII domain boundary of the KaiC hexamer. Furthermore, the model addresses the physical basis for temperature compensation concerning both the oscillation period and ATPase activity. Thus, the molecular clockwork perspective provides a framework for understanding the atomic design behind collective oscillations.

## INTRODUCTION

When three cyanobacterial proteins, KaiA, KaiB, and KaiC, are incubated with ATP in vitro, the phosphorylation level of the KaiC hexamer shows stable oscillations with a period of approximately 24 hours [1]. A proposed mechanism for these oscillations suggests that each KaiC hexamer acts like a clock, determining the rhythm at the single-molecule level [2, 3]. In this study, we explore this molecular clockwork hypothesis by developing a theoretical model of molecular oscillations.

KaiC is composed of two tandemly connected domains: CI and CII. The CII domain hydrolyzes a few ATP molecules per day for each monomer [4, 5]. The phosphate group produced from ATP hydrolysis in CII may be used to phosphorylate two amino-acid residues within the CII domain, Ser431 (S) and Thr432 (T). In contrast, the CI domain hydrolyzes approximately 10 ~ 12 ATP molecules per day for each monomer [4, 5]. This ATP consumption in CI has attracted significant interest, with both experimental [5–8] and theoretical [9, 10] studies revealing correlations between CI-ATPase reactions, the structural states of KaiC, and its phosphorylation levels. However, the reason why the oscillations of KaiABC require ATP consumption in CI remains unclear.

Takao Kondo addressed this issue by proposing that the CI-ATPase reactions determine the rhythm within individual KaiC hexamers [2, 3]. He suggested that there is feedback coupling between ATPase reactions and structural deformations in CI, which may lead to oscillations in CI’s structure. In turn, these oscillations can induce structural oscillations in CII through allosteric communication between the two domains. Such structural changes are expected to regulate the phosphorylation-dephosphorylation reactions in CII. Therefore, we can think of the structural oscillations driven by the CI-ATPase reactions as a “pendulum” or “pacemaker” that regulates KaiC’s phosphorylation oscillations. Furthermore, phosphorylation oscillations in CII should influence CI’s structure, thereby enhancing CI’s structural oscillations. Using the analogy of clockwork, Kondo referred to this feedback mechanism as “escapement” [2, 3] (Fig. 1A).

**FIG. 1.**
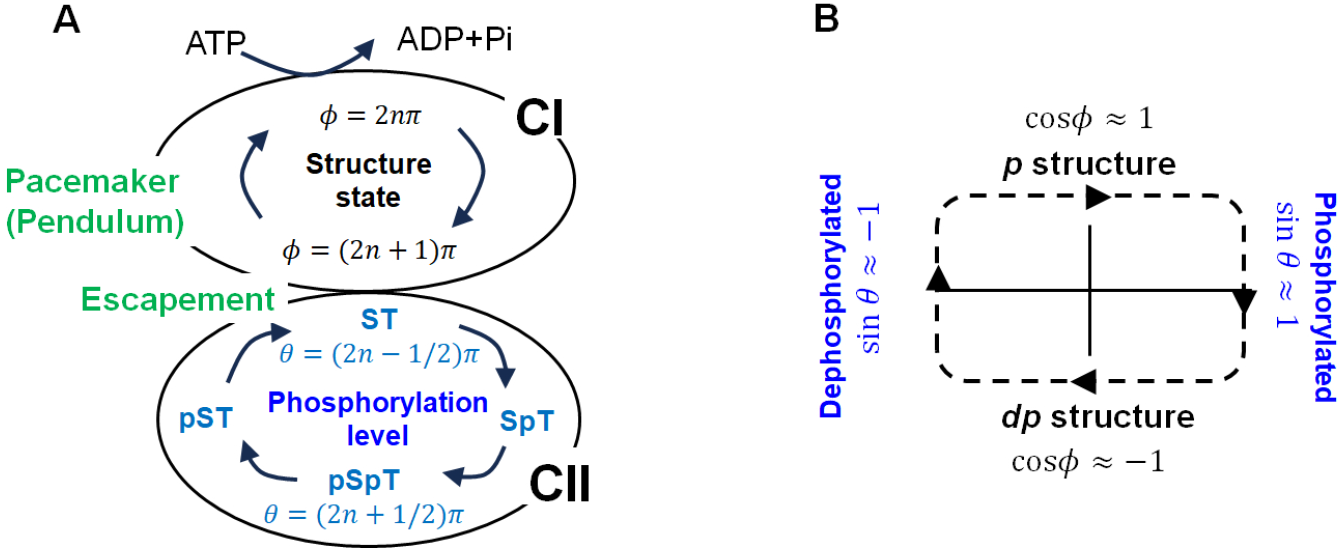
Molecular clockwork hypothesis. (**A**) The KaiC hexamer exhibits stochastic structural and chemical oscillations at the single molecular level. The structural oscillations, which function as a “pacemaker” or “pendulum,” are driven by the ATPase activity in CI. The phosphorylation level in CII oscillates through an “escapement” coupling with the pacemaker. We use phase variables to represent these oscillations: *ϕ* for the structural oscillations and *θ* for the phosphorylation oscillations. (**B**) Stable oscillations are achievable when *ϕ* and *θ* are coordinated. The *p* structure at *ϕ* ≈ 2*nπ* (*n* is an integer) facilitates phosphorylation, while the *dp* structure at *ϕ* ≈ (2*n* + 1)*π* allows for dephosphorylation. The phosphorylation states at *θ* ≈ (2*n*−1*/*2)*π*, in which both Ser431 (S) and Thr432 (T) are dephosphorylated, promotes transition from *dp* to *p* structures. Conversely, the state at *θ* ≈ (2*n* + 1*/*2)*π*, in which both S and T are phosphorylated (pSpT), destabilizes the *p* structure, turning it to the *dp* structure.

This molecular clockwork hypothesis requires testing by analyzing how atomic perturbations modify the rhythm. Specifically, we need to clarify the mechanism behind the significant changes in the oscillation period that occur when substituting an amino acid near the boundary between CI and CII [8]. Additionally, it is important to address the weak dependence of the oscillation period on temperature [1], a phenomenon known as temperature compensation. We also need to explain the experimentally observed correlation between oscillation frequency (1/period) and CI-ATPase activity across various mutations in KaiC [4, 5, 8]. Furthermore, the temperature compensation observed in ATPase activity [4, 8] also requires clarification. In this study, we propose a simple theoretical model to describe the coupled structural and chemical dynamics. We will examine whether the molecular clockwork hypothesis can account for the observed changes in oscillation period and temperature compensation, and analyze the role of CI-ATPase reactions in regulating these oscillations.

Another important aspect of the molecular clockwork hypothesis is the mechanism of synchronization. To generate the oscillations observed in a macroscopic test-tube sample, a large number of KaiC hexamers need to synchronize with one another. Various theoretical models have been developed to address this issue, proposing different hypotheses about how synchronization occurs. The earliest theoretical attempt suggested that direct interactions among multiple KaiC hexamers could lead to synchronization [11]. However, there is no experimental evidence for such direct interactions in the KaiABC system. Another proposed mechanism is the exchange of KaiC monomers among KaiC hexamers [12–15]. This exchange has indeed been observed in vitro [12, 16], suggesting it may play a role in synchronization.

The other important mechanism is the sequestration of KaiA molecules [10, 17–24]. When a significant amount of KaiA binds to specific states of KaiC, the concentration of free, unbound KaiA decreases. As a result, the rate of KaiA-assisted phosphorylation declines across many KaiC hexamers simultaneously, promoting synchronization among them. Experimental observations confirmed this KaiA sequestration mechanism, as the macroscopic oscillations disappear when an excess of KaiA molecules prevents effective sequestration [25]. Various theoretical models suggest different states of KaiC that bind and sequester KaiA: Some models highlight the lowly phosphorylated states during the phosphorylation phase [18, 19], while others focus on the highly phosphorylated states during the phosphorylation phase [17], or the lowly phosphorylated states during the dephosphorylation phase [20, 21].

Another proposed sequestration state is the KaiA-KaiB-KaiC complex [10, 22–24]. During the dephosphorylation phase, six KaiB molecules bind to the CI region of the six subunits of the KaiC hexamer. Electron microscopy observations revealed that a KaiA dimer binds to each KaiB molecule, enabling the incorporation of six KaiA dimers into the complex [26]. This complex, which forms during dephosphorylation, provides a clear platform for KaiA sequestration. Furthermore, this sequestration within the KaiA-KaiB-KaiC complex aligns with the experimentally observed phase entrainment in oscillations [23]. Therefore, in this study, we adopt this straightforward mechanism to explain the synchronization of KaiC hexamers.

This paper is organized as follows: First, we introduce phase variables that describe the phosphorylation levels and structural states in a KaiC hexamer. Next, we examine KaiA sequestration to demonstrate how KaiA globally couples KaiC hexamers through its sequestration. We then investigate the molecular clockwork hypothesis by analyzing the mean-field behaviors of an ensemble of molecular clocks. The mean-field analyses indicate that the interactions between the pacemaker and escapement processes within each KaiC hexamer determine the period of the ensemble-level oscillations. This mechanism for period determination aligns with the experimentally observed dependence of the oscillation period on perturbations, such as changes in temperature and various mutations. Finally, we analyze entropy production within this mean-field model to investigate the role of ATP consumption in the CI domain in regulating the oscillations.

## METHODS

### Phase model of molecular oscillators

We develop a phase model of the molecular clockwork hypothesis (Fig. 1). Observations from X-ray diffraction [27], biochemical analyses [28], NMR [29], and electron microscopy [30] have shown that the KaiC hexamer undergoes allosteric transitions between two structures. One structure can bind a KaiA dimer on the CII side, while the other can bind six KaiB monomers on the CI side. The structure that binds KaiA promotes phosphorylation in CII, which we refer to as the *p* structure. The structure that binds KaiB promotes dephosphorylation, which we refer to as the *dp* structure.

We hypothesize that stochastic CI-ATPase reactions trigger transitions between these two structures. Furthermore, when the structure influences ATPase-reaction rates, a feedback relationship may develop between them. Each structural transition is expected to occur within seconds, similar to other typical protein transitions. However, this feedback could bias these transitions, possibly resulting in limit-cycle oscillations with a period of hours [9, 10, 22, 23]. We assume the existence of such intramolecular cyclic behavior, as proposed by Kondo in his clockwork hypothesis [2, 3], and use this assumption as our starting point to test the hypothesis. For a possible model showing these limit-cycle oscillations, see Appendix A.

We define the phase variable *ϕ*_*i*_ as cos 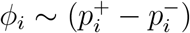 to indicate the position of the *i*th KaiC hexamer along the cycle. Here, 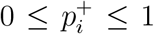 represents the probability that the *i*th KaiC hexamer adopts the *p* structure and 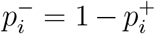 is the probability of adopting the *dp* structure. For assessing the coherence of the ensemble-level oscillations, we define the order parameter *r*_*ϕ*_ by

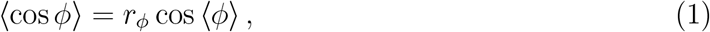

where 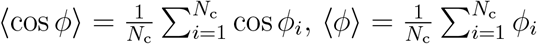, and *N*_c_ is the total copy number of KaiC hexamers in the solution.

Each CII domain has two sites, S and T, that can be phosphorylated. Therefore, a KaiC hexamer contains a total of 12 sites to be phosphorylated. We denote the number of phosphorylated sites as 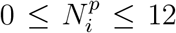 and define the phase variable *θ*_*i*_ of the *i*th KaiC hexamer as 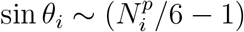. We also define *r*_*θ*_ by ⟨cos *θ*⟩ = *r*_*θ*_ cos ⟨*θ*⟩.

A coupling between the structural and phosphorylation states has been observed in various experiments [27–30]. In our present model, this coupling, referred to as the escapement in Fig. 1A, indicates *ϕ*_*i*_ ≈ *θ*_*i*_ during stable oscillations (Fig. 1B). An example cycle of oscillations proceeds as follows: When cos *ϕ*_*i*_ > 0 with *ϕ*_*i*_ ≈ 2*nπ* and *n* being an integer, the *p* structure predominantly appears, leading to the phosphorylation of S and T. This process results in a change of *θ*_*i*_ from (2*n* − 1*/*2)*π* to (2*n* + 1*/*2)*π*. Then, the charge distribution in the highly phosphorylated states at *θ*_*i*_ ≈ (2*n* + 1*/*2)*π* destabilizes the *p* structure, which drives a structural transition from *ϕ*_*i*_ ≈ 2*nπ* to *ϕ*_*i*_ ≈ (2*n* + 1)*π*, resulting in cos *ϕ*_*i*_ *<* 0. When cos *ϕ*_*i*_ *<* 0, the *dp* structure facilitates the dephosphorylation of pS and pT, causing *θ*_*i*_ to shift from (2*n*+1*/*2)*π* to (2*n*+3*/*2)*π*. Then, the dephosphorylated state at *θ*_*i*_ ≈ (2*n*+3*/*2)*π* makes the *dp* structure unstable, promoting a transition from *ϕ*_*i*_ ≈ (2*n* + 1)*π* to *ϕ*_*i*_ ≈ 2(*n* + 1)*π*, which returns the KaiC hexamer to the *p* structure and initiates the phosphorylation process for the next cycle.

This cycle operates on a timescale of hours, which is a process far from equilibrium. However, over shorter timescales of seconds, the system should be effectively in equilibrium. In this quasi-equilibrium, we assume that the free energy *U*_*θϕ*_ ∝ − cos(*θ* − *ϕ*) represents the escapement mechanism for phase alignment *ϕ* ≈ *θ*. With this expression, we expect that the time evolution of *θ*_*i*_ and *ϕ*_*i*_ is driven by forces proportional to −*∂U*_*θϕ*_*/∂θ*_*i*_ and −*∂U*_*θϕ*_*/∂ϕ*_*i*_, respectively. Therefore, using constants *b*_*θ*_ > 0 and *b*_*ϕ*_ > 0, the escapement coupling of *θ*_*i*_ and *ϕ*_*i*_ is modeled as

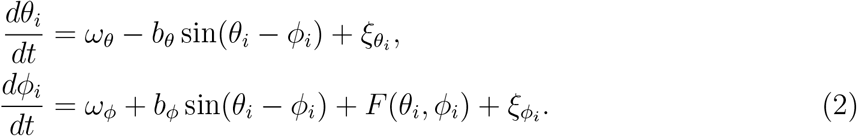

Here, the coupling *b*_*θ*_ should arise from the allosteric communication from CI to CII. We propose that the value of *b*_*θ*_ will decrease if the steric packing at the CI-CII boundary is loosened, while the coupling *b*_*ϕ*_, which reflects the effect of the CII-phosphorylation on CI, is determined by the charge distribution within CII, making it less sensitive to packing tightness. In the subsections “Mutations at the CI-CII domain boundary” and “Temperature compensation,” we will illustrate how this difference is critical for explaining the KaiC oscillations using our model.

*ω*_*θ*_ and *ω*_*ϕ*_ in Eq. 2 represent driving forces fueled by the CII-ATPase and CI-ATPase reactions, respectively. As the CI-ATPase is more active than the CII-ATPase, we consider *ω*_*ϕ*_ ≫ *ω*_*θ*_. Thus, *ω*_*ϕ*_ acts as the pacemaker, as illustrated in Fig. 1. 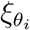 and 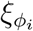 are Gaussian noises satisfying 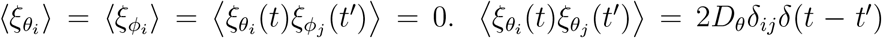 and 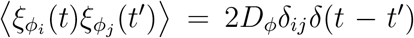, where *D*_*θ*_ and *D*_*ϕ*_ are constants representing the noise strength.

*F* (*θ*_*i*_, *ϕ*_*i*_) in Eq. 2 describes the effect of binding of KaiA to CII. Here, the probability to bind KaiA on KaiC is proportional to *a*_0_, which is the concentration of free, unbound KaiA dimers in the solution. To derive the expression for *a*_0_, we define total concentrations of KaiC hexamers and KaiA dimers in the solution as *c* and *a*, respectively. We also define the dissociation constant for a KaiA dimer from a KaiB subunit of the KaiB-KaiC complex as *K*. Then, we obtain the expression,

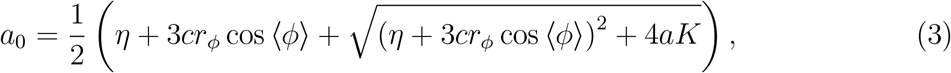

where *η* = *a* − 3*c* − *K* (Appendix B). Eq. 3 indicates that *a*_0_ shows oscillations as the phase ⟨*ϕ*⟩ proceeds. Using the constants *γ*_0_ and *γ*_1_, we approximate these oscillations as *a*_0_ ≈ *γ*_0_ + *γ*_1_*r*_*ϕ*_ cos ⟨*ϕ*⟩. See Appendix B and Fig.6 for the validy of this approximation. The oscillation strength *γ*_1_ is small when *η* + 3*cr*_*ϕ*_ cos ⟨*ϕ*⟩ ≤ *η* + 3*c <* 0. On the other hand, when the non-oscillatory amount *γ*_0_ is large, we have *η* + 3*cr*_*ϕ*_ cos ⟨*ϕ*⟩ ≈ *η* > 0. Therefore, the *a*_0_ oscillations are significant only when −3*c* ≲ *η* ≲ 0, or

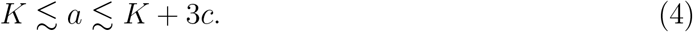

As demonstrated in the derivation of Eq. 3 in Appendix B, the *a*_0_ oscillations arise from the sequestration of KaiA to the KaiA-KaiB-KaiC complex. Thus, Eq. 4 serves as a necessary condition for the effective sequestration.

When a KaiA dimer binds to CII, it stabilizes the *p* structure while destabilizing the *dp* structure. As the probability of a KaiA dimer binding to CII is proportional to *a*_0_, we represnt this structural effect by the quasi-equilibrium free energy 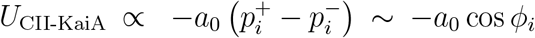. Then, the driving force for phase progress is proportional to −*∂U*_CII-KaiA_*/∂ϕ*_*i*_, and the force can be further enhanced when the structural state and phosphorylation level are in alignment without frustration [31], which is an aspect of the escapement mechanism. Therefore, using a constant *β*_0_ > 0, we propose that the function *F* (*θ*_*i*_, *ϕ*_*i*_) has the form,

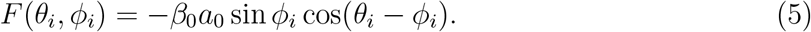

Here, we focus on slow relaxation or fluctuations that occur over multiple oscillation periods. In this case, it is essential to retain the slow-varying term in Eq. 5, as demonstrated in the derivation of Kuramoto models for chemical reactions [32]. When the oscillation amplitude is large with *r*_*ϕ*_ ≈ 1 in Eq. 1, we express the phase as *ϕ*_*i*_ = *y*_*i*_ + *ωt*, with ⟨*y*_*i*_⟩ = 0 and ⟨*ϕ*_*i*_⟩ = ⟨*ϕ*⟩ = *ωt*. Then, Eq. 5 leads to

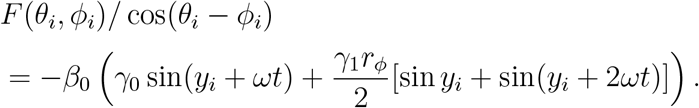

In this expression, the slow term is proportional to 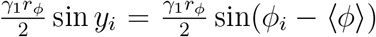. When *ω*_*ϕ*_*/β*_0_ is sufficiently large, we can disregard the fast-varying terms proportional to *γ*_0_ sin(*y*_*i*_ + *ωt*) and 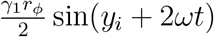. Thus, we substitute Eq. 5 with the expression:

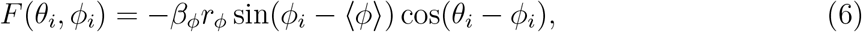

with *β*_*ϕ*_ = *β*_0_*γ*_1_*/*2. Eq. 6 represents the scenario that the molecular oscillators {*θ*_*i*_, *ϕ*_*i*_} synchronize through the global coupling of sin(*ϕ*_*i*_ − ⟨*ϕ*⟩). This global coupling occurs as KaiA molecules diffuse globally throughout the solution.

The effectiveness of KaiA sequestration in generating the global coupling can be demonstrated by numerically calculating Eqs. 2–3 with Eq. 5 for the system containing *N*_c_ copies of KaiC hexamers. Fig. 2 illustrates examples of the calculated oscillations. It shows that the amplitude of the simulated ensemble-level oscillations, which approximately corresponds to *r*_*ϕ*_, is significant only when the KaiA concentration is within a specific range. This calculated range is narrower than and slightly shifted from what was suggested by the rough estimation of Eq. 4. When the KaiA concentration falls outside this range, the sequestration becomes ineffective, thereby reducing *r*_*ϕ*_. This result aligns with the experimental observations, showing that the macroscopic oscillations exist only within a specific range of KaiA concentration, 0.3 ≲ *a/c*_0_ ≲ 4, when *c* = *c*_0_ ≈ 0.6 *µ*M [25]. In the following, we use Eq. 6 instead of Eq. 5, assuming that the KaiA concentration is within the range that supports oscillations.

**FIG. 2.**
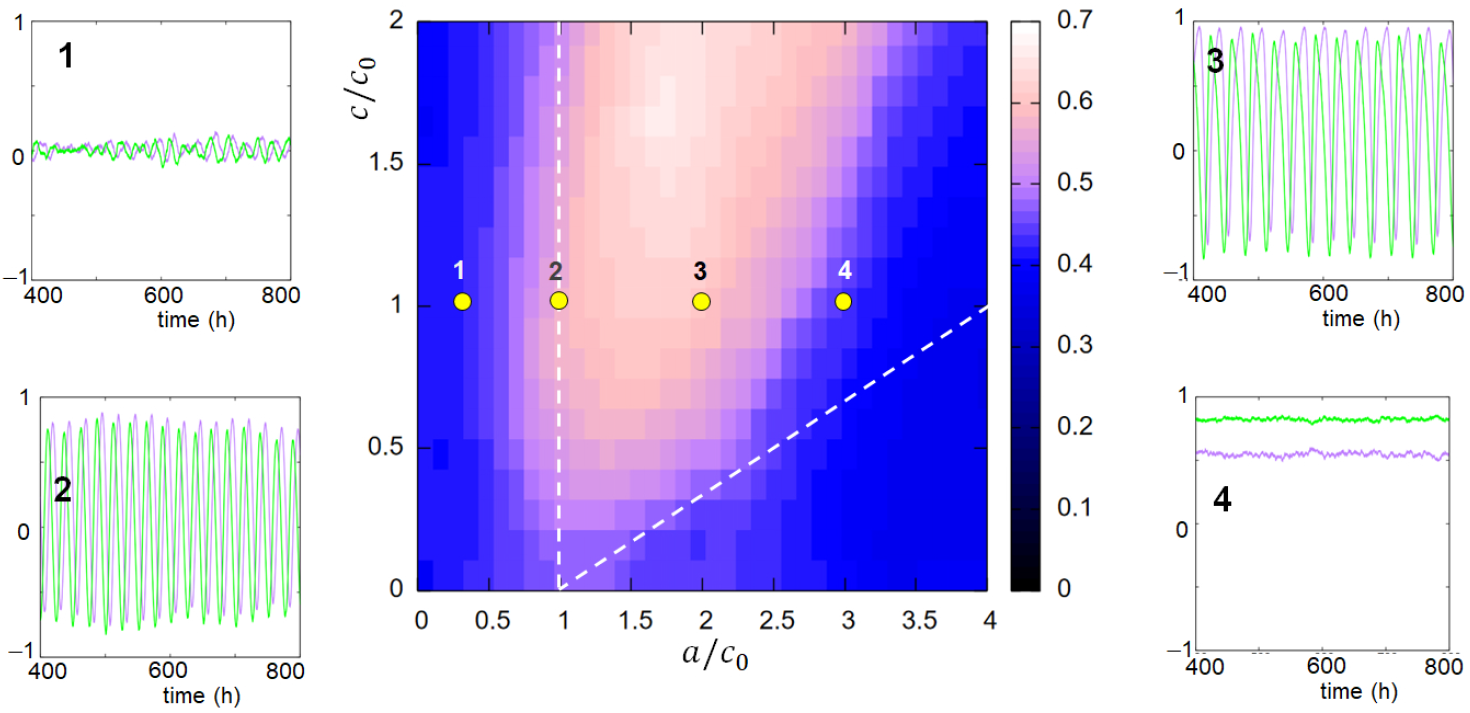
Synchronization of KaiC oscillations through sequestration of KaiA to the KaiA-KaiB-KaiC complex. The phase diagram of oscillations is shown by plotting the amplitude of temporal variation of the ensemble-level phosphorylation. The amplitude is plotted by a color map on a plane defined by the normalized total concentrations of KaiA dimers, *a/c*_0_, and KaiC hexamers, *c/c*_0_, where *c*_0_ is the total concentration of KaiC hexamers in the standard solution. The KaiA sequestration is effective only within a specific range of *a*, making the amplitude large in that area. The area between two dashed lines is the region satisfying the condition of Eq. 4. Also shown are trajectories of temporal evolution of the ensemble-level oscillations of structural state (green) and phorphorylation level (purple) at numbered points on the plane. Calculated using Eqs. 2–3 and Eq. 5, with *N*_c_ = 100, *K/c*_0_ = 1, *ω*_*θ*_ = 0, *ω*_*ϕ*_ = 2*π/*22 */*h, *b*_*θ*_ = 0.1 */*h, *b*_*ϕ*_ = 1 */*h, *β*_0_ = 2 */*h, and *D*_*ϕ*_ = *D*_*θ*_ = 0.01 */*h.

### Mean-field approximation

We define the distribution of molecular oscillators, *P* ({*θ*_*i*_, *ϕ*_*i*_}), satisfying the following equation, which is equivalent to Eq. 2,

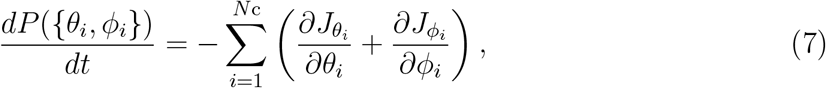

where 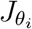 and 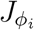 are probability currents,

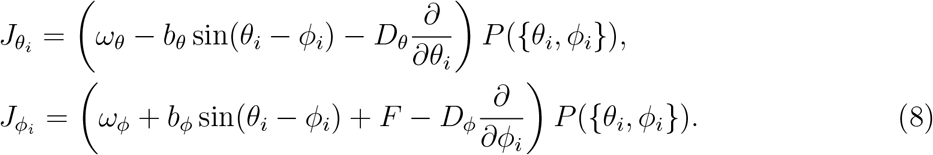

The global coupling of molecular oscillators described in Eq. 6 suggests that the mean-field approximation is a straightforward approach. In this approximation, the distribution is factorized as 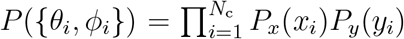 with *x*_*i*_ = *θ*_*i*_ − *ϕ*_*i*_ and *y*_*i*_ = *ϕ*_*i*_ − *ωt*, where *x*_*i*_ and *y*_*i*_ are variables representing fluctuations around coherent oscillations, satisfying ⟨*y*_*i*_⟩ = 0 and ⟨*ϕ*_*i*_⟩ = *ωt*. Here, *ω* is the frequency of the ensemble-level oscillations. By employing Eq. 6 along with this factorization in Eq. 7, we obtain the equations for *P*_*x*_ and *P*_*y*_. These equations can then be solved to yield a solution representing the steady distributions of *x* and *y* (Appendix C).

With this steady-state solution, the amplitude of the ensemble-level oscillations is *r*_*ϕ*_ = ⟨cos *y*⟩. This expression leads to a self-consistent relation for *r*_*ϕ*_ (Appendix D), predicting the critical emergence of oscillations as the couplings *b*_*θ*_, *b*_*ϕ*_, or *β*_*ϕ*_ increase. Specifically, by using 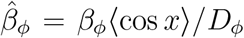, we find *r*_*ϕ*_ = 0 for 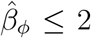, and 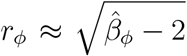 for 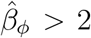. For a large 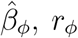 approaches to 1. This mean-field prediction aligns well with the numerical solutions obtained from Eqs. 2 and 6 for the system containing *N*_c_ copies of KaiC hexamers, as illustrated in the phase diagram (Fig. 3A) and in the oscillation amplitudes (Fig. 3B).

**FIG. 3.**
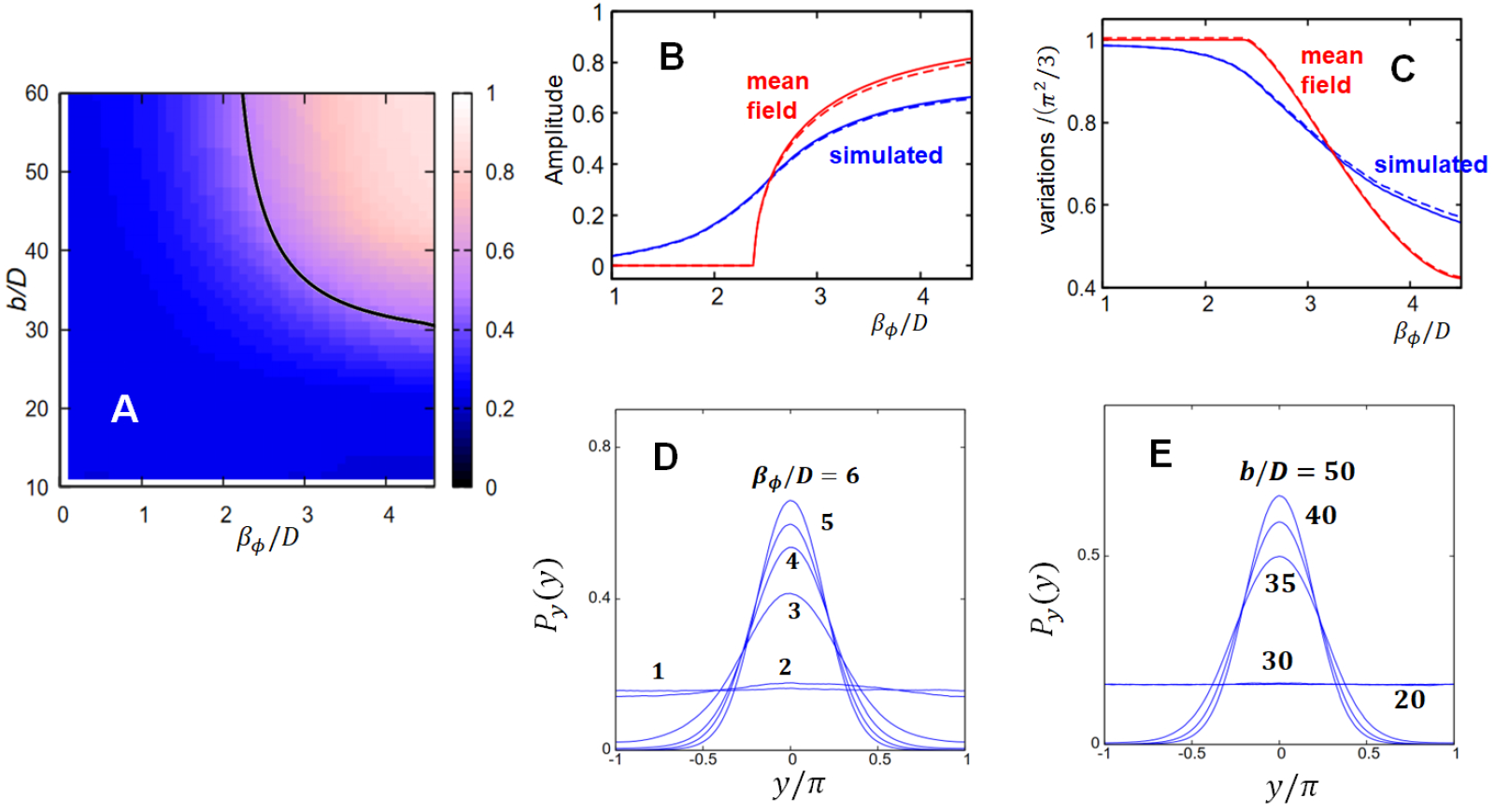
The ensemble-level oscillations numerically calculated with the phase model (Eqs. 2 and 6) of *N*_c_ copies of KaiC hexamers are compared with the corresponding mean-field results. (**A**) The phase diagram of oscillations is shown on a plane defined by the normalized intermolecular coupling *β*_*ϕ*_*/D*_*ϕ*_ = *β*_*ϕ*_*/D*, and the normalized intramolecular coupling *b*_*θ*_*/D*_*θ*_ = *b*_*ϕ*_*/D*_*ϕ*_ = *b/D*. Amplitude of the temporal variation of the ensemble-level phosphorylation, numerically calculated with the phase model, is plotted by a color map. The black line is the mean-field critical line defined by 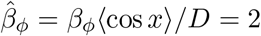. (**B**) Amplitudes of the temporal variation of the ensemble-level structural state (*r*_*ϕ*_, blue, real line) and phosphorylation level (*r*_*θ*_, blue, dashed line), numerically calculated with the phase model, are plotted as functions of the intermolecular coupling strength, *β*_*ϕ*_*/D*, and compared with the corresponding amplitudes obtained from the mean-field theory for *r*_*ϕ*_ (red, real line) and *r*_*θ*_ (red, dashed line). Here, *r*_*ϕ*_ is defined by Eq. 1, and *r*_*θ*_ is defined as ⟨cos *θ*⟩ = *r*_*θ*_ cos ⟨*θ*⟩. (**C**) Variances of oscillations numerically calculated with the phase model, for the structural state (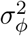, blue, real line) and the phosphorylation level (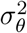, blue, dashed line), are compared with the corresponding variances obtained from the mean-field theory for 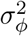 (red, real line) and 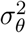 (red, dashed line). (**D, E**) The distribution of oscillations numerically calculated with the phase model, *P*_*y*_(*y*), shows a sudden change at around the critical line predicted by the mean-field theory, where *y* = *ϕ* − ⟨*ϕ*⟩ represents fluctuations around the coherent oscillations. The parameters used are *K/c*_0_ = 1, *ω*_*θ*_ = 0, *ω*_*ϕ*_ = 2*π/*12 */*h, and *D*_*ϕ*_ = *D*_*θ*_ = *D* = 0.01 */*h for **A**–**E**, *b/D* = 50 for **B**–**D**, and *β*_*ϕ*_*/D* = 6 for **E**. In the numerical calculations with the phase model using Eqs. 2 and 6, we assumed *N*_c_ = 100 in **A**–**C**, and *N*_c_ = 1000 in **D** and **E**. The mean-field results were obtained by numerically solving the self-consistent relation (Appendix C).

Variances of fluctuations across the ensemble of KaiC hexamers, 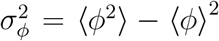 and 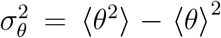, are also calculated using the mean-field approximation (Appendix E):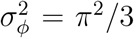 when the coupling is small for 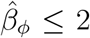, which is reduced to 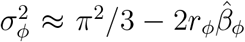 when 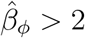 with *r*_*ϕ*_ > 0. When the coupling 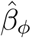 is large, we have

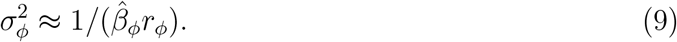

We have 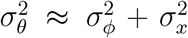, where 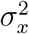 remains small when *b*_*θ*_ and *b*_*ϕ*_ are sufficiently large. These mean-field results also align with the numerical solutions obtained from Eqs. 2 and 6 (Fig. 3C). Additionally, distribution *P*_*y*_(*y*), numerically derived from Eqs. 2 and 6, shows a sudden change at around the critical value of the coupling strengths predicted by the mean-field approximation (Fig. 3D and Fig. 3E). Thus, the mean-field approximation effectively captures the behavior of the phase model defined by Eqs. 2 and 6.

Dropping the index *i* from the mean-field equations, the Langevin equations for *x* and *y* are

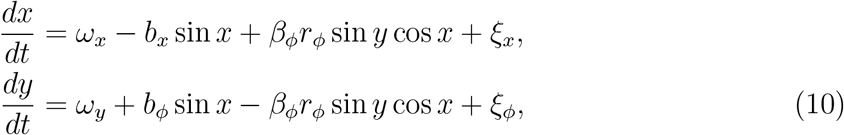

where *ω*_*x*_ = *ω*_*θ*_ − *ω*_*ϕ*_, *ω*_*y*_ = *ω*_*ϕ*_ − *ω, b*_*x*_ = *b*_*θ*_ + *b*_*ϕ*_, and *ξ*_*x*_ = *ξ*_*θ*_ − *ξ*_*ϕ*_. When coherent oscillations persist, the fluctuations *x* and *y* around these oscillations satisfy ⟨*dx/dt*⟩ ≈ ⟨*dy/dt*⟩ ≈ 0. In this scenario, after taking the average in Eq. 10 and using the mean-field relation of ⟨sin *y* cos *x*⟩ = ⟨sin *y*⟩ ⟨cos *x*⟩ = 0, we have 0 = *ω*_*x*_ − *b*_*x*_ ⟨sin *x*⟩ and 0 = *ω*_*y*_ + *b*_*ϕ*_ ⟨sin *x*⟩, which lead to

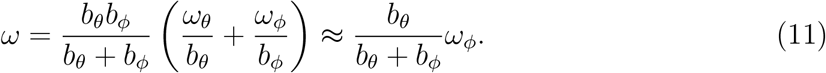

In Eq. 11, we used *ω*_*ϕ*_ ≫ *ω*_*θ*_. Thus, the bare value of the pacemaker frequency, *ω*_*ϕ*_, is renormalzed by the escapement coupling through *b*_*θ*_ and *b*_*ϕ*_ to give rise to the frequency *ω* of the ensemble-level oscillations.

### Parameters

The mean-field analyses highlight the importance of distinguishing among three regimes based on coupling strength: weak 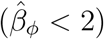, strong 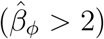, and critical 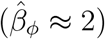 coupling regimes. When this distinction is clear under a given condition, many of the model’s mean-field predictions become independent of particular parameter choices in that condition. For example, Eq. 11 in the strong coupling regime illustrates the relationship between *ω/ω*_*ϕ*_ and *b*_*θ*_*/b*_*ϕ*_. As long as the system remains within the strong coupling regime, this relationship is valid regardless of the exact values of other parameters such as the noise strengths *D*_*θ*_ and *D*_*ϕ*_, the concentration of KaiA dimers *a*, the concentration of KaiC hexamers *c*, or the dissociation constant *K*. However, to quantitatively compare these mean-field results with experimental observations, it is helpful to assume specific values for the parameters *ω*_*ϕ*_, *ω*_*θ*_, *b*_*θ*_, and *b*_*ϕ*_. In this subsection, we will discuss these parameters in detail.

We begin by proposing that *ω*_*ϕ*_ ≫ *ω*_*θ*_. In our hypothesis, the structural oscillations of *ϕ*, which are driven by ATP consumption in CI, act as the primary pacemaker of the KaiC oscillations. In contrast, the phosphorylation-level oscillations in CII, represented by *θ*, are subsidiary to the oscillations of *ϕ*. To reflect this relationship, we assume the condition *ω*_*ϕ*_ ≫ *ω*_*θ*_, which allows us to neglect the effects of *ω*_*θ*_ in many cases. While direct experimental confirmation of this inequality has not yet been achieved, our assumption is consistent with experimental observations indicating that the ATPase rate is significantly higher in CI than in CII. Therefore, we consider this assumption to be fundamental within the framework of the molecular clockwork hypothesis.

We next estimate *ω*_*ϕ*_ by comparing it with experimental data from the truncated CI. When the CII domain is disconnected from the CI, the remaining CI complex still exhibits ATPase activity [5, 33]. In our model, this truncated CI is characterized by the conditions where *b*_*θ*_ = *b*_*ϕ*_ = *ω*_*θ*_ = 0. Therefore, we can estimate *ω*_*ϕ*_ based on the temperature dependence of the ATPase activity of the truncated CI. The ATPase activity is quantified by the number of ADP molecules released over a 24-hour period for a truncated CI subunit at a temperature *T*, which we denote as *N* ^tCI−ATP^(*T*). Experimental data show that *N* ^tCI−ATP^(*T*) follows a linear trend in an Arrhenius plot, and the ratio 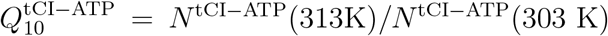 is approximately 1.53 [33]. This value of 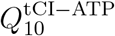, along with a relationship that will be explained later in Eq. 23, leads to the expression of 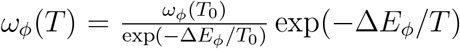 with the activation energy Δ*E*_*ϕ*_ ≈ 7.5*k*_B_*T*_0_ at around *T*_0_ = 300 K and *k*_B_ being the Boltzmann constant. This expression will be used in the subsections “Mutations at the CI-CII domain boundary” and “Temperature compensation.”

The couplings between *θ* and *ϕ, b*_*θ*_ and *b*_*ϕ*_ in our model, play an important role in the molecular clockwork hypothesis. Although direct measurements of *b*_*θ*_ and *b*_*ϕ*_ have not yet been conducted, we estimate these values based on physical arguments regarding protein conformation dynamics [34–36]. Frauenfelder demonstrated, through the dielectric relaxation spectra of myoglobin, that the dynamics of protein conformation are viscous, primarily due to the influence of solvent molecules surrounding the protein [34]. The observed kinetic rate of the conformational relaxation is given by

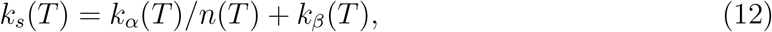

where *k*_*α*_(*T*) reflects the *α*-relaxation of the bulk solution across a broad temperature range, following the Vogel-Fulcher relaxation law *k*_*α*_(*T*) ~ exp(−*T*_2_*/*(*T* − *T*_1_)) [34–36]. The term *k*_*β*_ corresponds to the solid-like *β*-relaxation of the protein, with *k*_*α*_(*T*)*/n*(*T*) ≫ *k*_*β*_(*T*) at *T* ≈ *T*_0_ [34]. Here, *n*(*T*) ≫ 1 indicates how protein responds to solvent dynamics [35]. We expect that the fast relaxation, characterized by a larger value of *k*_*s*_, occurs when two domains are weakly coupled, resulting in large conformational fluctuations. Therefore, by neglecting a small parameter *k*_*β*_ in Eq. 12, we propose that *b*_*θ*_ is inversely related to *k*_*s*_ as

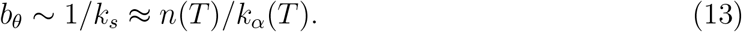

Additionally, the random first-order transition theory of glassy dynamics suggests that *n*(*T*) decreases as the protein conformation becomes less constrained, showing larger fluctuations [35]. When a residue near the CI-CII domain boundary is replaced with a residue that has a smaller volume *v* than the wild-type residue with volume *v*_0_, the protein conformation should indeed be less constrained. To capture this trend while ensuring numerical stability in our calculations, we propose an exponentiated form of the relationship *n*(*T*) ∝ exp[*ϵ*(*T*)(*v/v*_0_ − 1)] with *ϵ*(*T*) > 0. Given that increasing temperature masks the steric packing effects, making them less apparent, we anticipate that *ϵ*(*T*) is a decreasing function of temperature *T*. Therefore, we suggest the approximation *ϵ*(*T*) ≈ *ϵ*_0_ + *ϵ*_1_(*T* − *T*_0_), with *ϵ*_0_ > 0 and *ϵ*_1_ *<* 0.

In contrast, *b*_*ϕ*_ reflects changes in the charge distribution at 12 sites in CII and is expected to be less sensitive to conformational fluctuations. Therefore, we assume that the effects of changes in *v* or *T* primarily manifest through changes in *b*_*θ*_. Thus, we propose the following relationship:

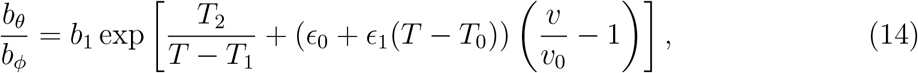

where *b*_1_ = *b*_0_ exp [−*T*_2_*/* (*T*_0_ − *T*_1_)]. We use this expression in the subsections “Mutations at the CI-CII domain boundary.” and “Temperature compensation.” From the data on myoglobin, we find *T*_2_ is approximately several hundred K, while *T*_1_ should be *T*_1_ *< T*_cross_, where *T*_cross_ ≈ 210 K is the cross-over temperature satisfying the condition *k*_*α*_(*T*_cross_)*/n*(*T*_cross_) ≈ *k*_*β*_(*T*_cross_) [34].

We believe that recognizing the importance of coupling parameters *b*_*θ*_ and *b*_*ϕ*_, as predicted by our model, will encourage experimental efforts to measure them. One potential method to assess *b*_*θ*_ and *b*_*ϕ*_ is to evaluate the following response functions in the non-oscillatory KaiC solutions, which do not contain KaiA or KaiB:

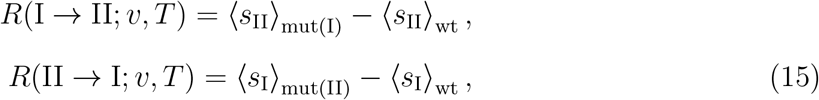

where *s*_I_ and *s*_II_ are any structural features, such as distances between monitoring sites, in the CI and CII domains, respectively. ⟨· · · ⟩_wt_ represents the average in wild-type KaiC, while ⟨· · · ⟩_mut(I)_ is the average in a mutant KaiC where an amino acid residue in the CI domain is replaced with one that does not alter the charge distribution. On the other hand, ⟨· · · ⟩_mut(II)_ denotes the average in a mutant KaiC in which an amino acid residue in the CII domain is substituted with one that modifies the charge distribution, simulating phosphorylation or dephosphorylation. The quantities *R*(I → II; *v, T*) and *R*(II → I; *v, T*) are expected to depend on the side-chain volume *v* of the residue at the CI-CII boundary, as well as the temperature *T*. Consequently, structural measurements that reveal the differences in how *R*(I → II; *v, T*) and *R*(II → I; *v, T*) depend on *v* and *T* —through methods such as molecular dynamics simulations, X-ray analyses, electron microscopy, or NMR measurements—should offer insights into the asymmetry between *b*_*θ*_ and *b*_*ϕ*_ in our model.

## RESULTS

### Mutations at the CI-CII domain boundary

The substitution of the residue Tyr402, located at the boundary between CI and CII, dramatically alters the period of the ensemble-level oscillations, changing it from 15 hours to 158 hours, where the oscillation frequency of the mutants correlates with the side-chain volume of the substituted residue [8]. We write *b*_*θ*_ and *b*_*ϕ*_ in the form of *b*_*θ*_ ∝ exp [*f*_*θ*_(*v, T*)] and *b*_*ϕ*_ ∝ exp [*f*_*ϕ*_(*v, T*)], where *f*_*θ*_(*v, T*) and *f*_*ϕ*_(*v, T*) are functions of temperature *T* and the volume *v* of the residue at the boundary of CI and CII.

Based on the assumptions discussed in the “Parameters” subsection, both *b*_*θ*_ and *b*_*ϕ*_ should be increasing functions of the volume *v* with *∂f*_*θ*_*/∂v* > 0 and *∂f*_*ϕ*_*/∂v* > 0. However, as the coupling from CI to CII, represented by *b*_*θ*_, should be through the steric packing at the domain boundary, we expect that *b*_*θ*_ depends more sensitively to *v* than *b*_*ϕ*_, which is represented as *∂*Δ*f/∂v* > 0 with Δ*f* = *f*_*θ*_ − *f*_*ϕ*_. A possible functional form of Δ*f* is presented in Eq. 14 as Δ*f* = *T*_2_*/*(*T* − *T*_1_) + *ϵ*(*T*)(*v/v*_0_ − 1) with *ϵ*(*T*) = *ϵ*_0_ + *ϵ*_1_(*T* − *T*_0_) > 0. Then, from Eq. 11, we obtain

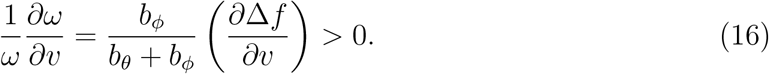

indicating that the frequency *ω* of the ensemble-level oscillations is an increasing function of the residue volume *v* at the CI-CII domain boundary.

In Fig. 4A, this mean-field prediction is compared with the experimentally observed periods of mutants [8]. Here, the oscillation period 2*π/ω* was calculated using Eqs. 11 and 14, with the parameters *b*_0_ and *ϵ*_0_ adjusted to fit the experimental data at *T* ≈ *T*_0_. Notably, even a moderate change in the ratio *v/v*_0_ results in nearly a tenfold change in the calculated period, which is consistent with the experimental observations. In this way, the molecular clockwork hypothesis explains the observed drastic mutational effects at position 402 in KaiC. The effects of varying parameters *b*_0_ or *ϵ*_0_ are also illustrated in Fig. 4A, suggesting that mutations at different sites should correspond to distinct parameter values, which lead to varying degrees of period modulation.

**FIG. 4.**
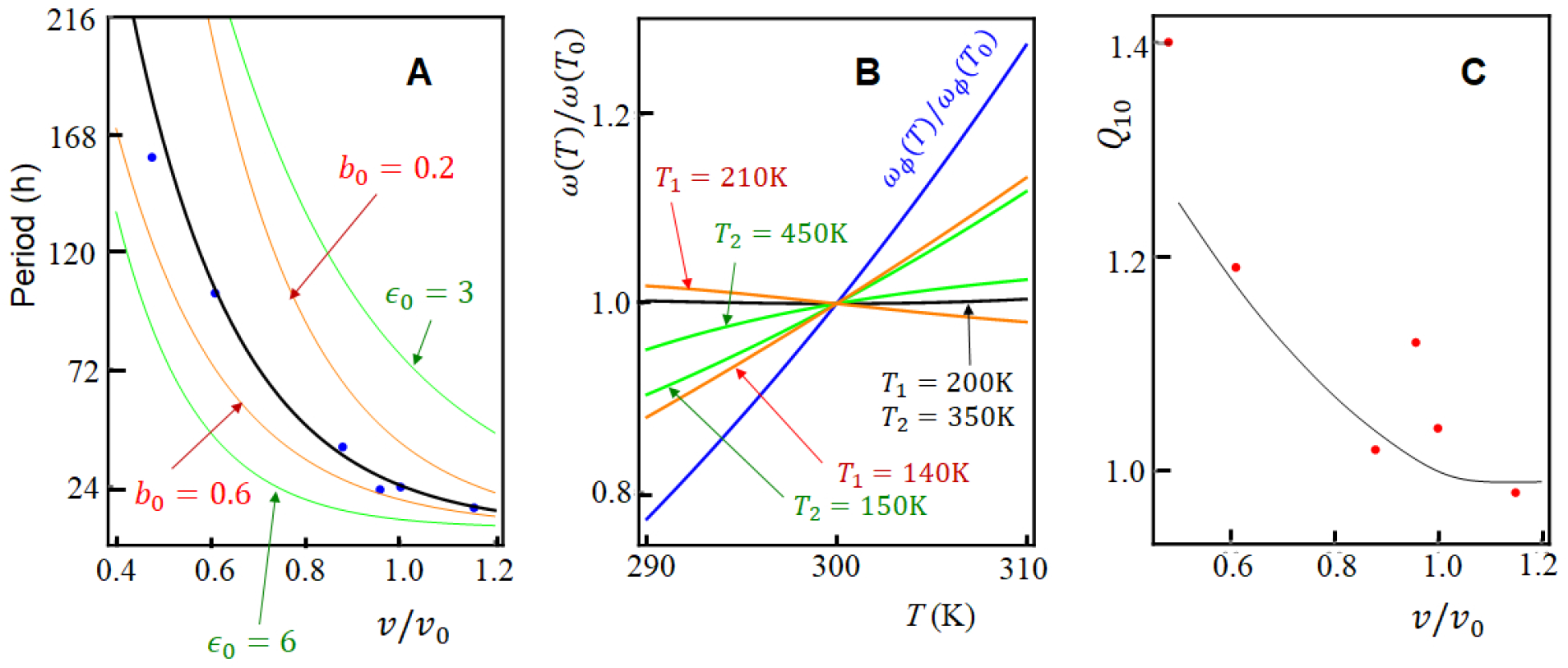
The bare frequency *ω*_*ϕ*_ of the structural oscillations is renormalized to yield the observed frequency *ω* through the escapement mechanism of the intramolecular couplings. Eqs. 11 and 14 are employed to calculate this renormalization effect. (**A**) The renormalization effect explains a significant change in the oscillation period 2*π/ω*. The dots represent the experimentally observed periods resulting from the substitution of residue 402 at the CI-CII boundary [8]. These periods are plotted as a function of *v/v*_0_, where *v* is the side-chain volume of the substituted residue and *v*_0_ represents the volume of the wild-type residue. The theoretical results, calculated with *ω*_*ϕ*_(*T*_0_) = 2*π/*7 h^−1^ at *T*_0_ = 300 K, are dranw by lines: the black line corresponds to *b*_0_ = 0.4 and *ϵ*_0_ = 4.4; the orange lines show *ϵ*_0_ = 4.4 with *b*_0_ either 0.2 or 0.6; and the green lines denote *b*_0_ = 0.4 with *ϵ*_0_ either 3 or 6. (**B**) The temperature dependence of the frequency is plotted as *ω*(*T*)*/ω*(*T*_0_) with *v/v*_0_ = 1 and *b*_0_ = 0.4. The black line is for *T*_1_ = 200 K and *T*_2_ = 350 K. For the orange lines, *T*_1_ is either 210 K or 140 K while *T*_2_ = 350 K. For the green lines, *T*_2_ = 150 K or 450 K with *T*_1_ = 200 K. Compared is the temperature dependence of the bare frequency of the pacemaker, *ω*_*ϕ*_(*T*) ∝ exp [−Δ*E/*(*k*_B_*T*)], with Δ*E* = 7.5*k*_B_*T*_0_ (blue line). (**C**) *Q*_10_ for the oscillation period. The red dots indicate the experimentally obtained *Q*_10_ values from mutants in which residue 402 is substituted [8]. The line represents the theoretical *Q*_10_ calculated with *ϵ*_1_ = −0.06 K^−1^ and the same parameters used for the black line in panel **B**.

### Temperature compensation

The frequency of KaiABC oscillations remains largely unaffected by the temperature change, showing temperature compensation. Temperature compensation is observed in many organisms as a hallmark of the stable circadian oscillations: With a typical biochemical activation energy of Δ*E* = 10 ~ 20*k*_B_*T*_0_ at *T*_0_ = 300 K, the reaction rate increases as *T* is increased from *T*_0_ to *T*_0_ + 10 K, resulting in a ratio of *Q*_10_ = (the rate at *T* = *T*_0_ + 10 K)*/*(the rate at *T* = *T*_0_) = 1.4 ~ 2. However, for many organisms, the frequency of circadian oscillations changes only slightly, with 0.9 ≲ *Q*_10_ ≲ 1.1 [37].

A possible explanation of temperature compensation in cells suggests that the inactivation or degradation of transcription factors is enhanced at higher temperatures to compensate for the acceleration of other reactions [38]. However, the factors within the KaiABC system do not exhibit such inactivation or degradation. Another hypothesis suggests that the temperature-dependent competition between KaiA binding to different states of KaiC mitigates the effects of temperature [19]. However, such temperature-dependent competition of binding affinity has yet to be experimentally validated. Therefore, we need to explore alternative scenarios to explain temperature compensation in the KaiABC system.

Based on the assumptions discussed in the “Parameters” subsection, we anticipate that the intramolecular couplings are weakened when conformational fluctuations of KaiC hexamer becomes large. We expect that the coupling from CI to CII, represented by *b*_*θ*_, is more sensitive to this effect than the coupling from CII to CI, represented by *b*_*ϕ*_. Therefore, we expect *∂*Δ*f/∂T <* 0. We consider that the bare frequency *ω*_*ϕ*_ depends on temperature as *ω*_*ϕ*_ ∝ exp [−Δ*E/*(*k*_B_*T*)]. Then, Eq. 11 leads to

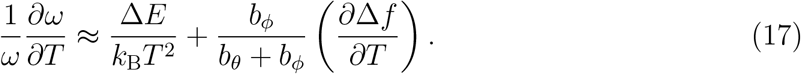

In this expression, with *∂*Δ*f/∂T <* 0, the first and second terms cancel each other to provide a small *∂ω/∂T*.

We anticipate that *b*_*θ*_ is correlated to the viscosity of domain motions, which may be inversely related to the relaxation rates in the protein’s conformation. The Vogel-Fulcher relation for glass-forming liquids could be an appropriate model for representing these rates [34]. Therefore, we use Eq. 14 for evaluating Δ*f*, with *T*_1_ = 200 K as estimated for the solvent effect on structural relaxation [34]. An example calculation for *ω* in Fig. 4B with *T*_2_ = 350 K and Δ*E* = 7.5 *k*_B_*T*_0_ yields *Q*_10_ = 1.01, indicating temperature compensation.

Dependence of *ω* on parameters *T*_1_ and *T*_2_ are shown in Fig. 4B. From Fig. 4B, we find that *Q*_10_ depends on both *T*_1_ and *T*_2_; for *T*_2_ = 350 K, we find *Q*_10_ = 0.98 (*T*_1_ = 210 K) and *Q*_10_ = 1.16 (*T*_1_ = 120 K), while for *T*_1_ = 200 K, *Q*_10_ = 1.03 (*T*_2_ = 450 K) and *Q*_10_ = 1.12 (*T*_2_ = 150 K). Since Eq. 12 suggests that the parameters *T*_1_ and *T*_2_ are determined by the relaxation rates of the bulk solvent, our model predicts that the degree of temperature compensation of oscillation period depends on the temperature dependence of viscosity of the bulk solution.

Various mutations at position 402 show that substitution to a residue with the smaller side-chain volume *v* leads to the larger *Q*_10_, as plotted in Fig. 4C [8]. In Fig. 4C, this dependence of *Q*_10_ on *v* is compared with the calculated *Q*_10_ for *T*_1_ = 200 K, *T*_2_ = 350 K and *ϵ*_1_ = −0.06 K^−1^. A correlation between the observe and calculated data found in Fig. 4C supports the consistency of Eqs. 11 and 14.

Thus, the molecular clockwork hypothesis effectively explains the physical mechanism of temperature compensation, as the coupling of the escapement, *b*_*θ*_ and *b*_*ϕ*_, and the pacemaker, *ω*_*ϕ*_, determines the rhythm.

### Dissipation and ATPase activity

The free energy gained by KaiC’s ATPase activity dissipates into the solution after one cycle of oscillations, which should be quantified as a house-keeping [39–41] or adiabatic [42–44] component of heat dissipation. In stable oscillations, we can write this adiabatic component as 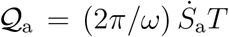, where 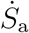 is the entropy production rate in the steadily oscillating state.

The concept of entropy production in the KaiABC system has been explored under the assumption of the monomer-exchange mechanism [15], as well as the KaiA sequestration mechanism [21] for synchronization. Furthermore, entropy production in a simplified model has been discussed, assuming the sequestration mechanism [45]. In this study, we demonstrate that the phase model offers a clear understanding of entropy production.

The entropy production rate per KaiC hexamer in a nonequilibrium steady state, whether in the non-oscillatory stationary state or the steadily oscillating state, has the expression [44] of 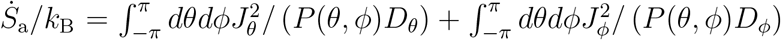. In the mean-field approximation, using *P* (*θ, ϕ*) = *P*_*x*_(*x*)*P*_*y*_(*y*), we can show that 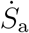 consists of several components (Appendix F): one that represents oscillations, and another that accounts for the intramolecular coupling with *b*_*θ*_ and *b*_*ϕ*_. The other component arises from global intermolecular coupling *β*_*ϕ*_*r*_*ϕ*_.

When the couplings are weak as *b*_*θ*_*/D*_*θ*_ ≪ 1, *b*_*ϕ*_*/D*_*ϕ*_ ≪ 1, and *β*_*ϕ*_*/D*_*ϕ*_ ≪ 1, the ensemble-level oscillations disappear with *r*_*ϕ*_ = 0, which eliminates the contribution from the global coupling. However, the *ϕ* and *θ* oscillations within individual molecules remain, yielding

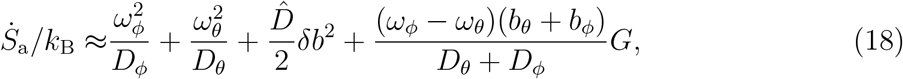

where 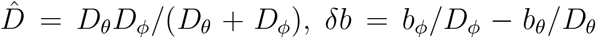, and *G* is a factor of *O*(*b*_*θ*_*/D*_*θ*_) and *O*(*b*_*ϕ*_*/D*_*ϕ*_) (Appendix F). The terms 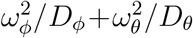 make a major contribution, representing the dissipation caused by the highly active *ϕ* oscillations of structure and the less active *θ* oscillations of phosphorylation. The term proportional to *δb*^2^ accounts for the intramolecular coupling between *ϕ* and *θ*, representing a “friction” between them. The last term arises because the more active movement of *ϕ* “drags” the less active movement of *θ*.

On the other hand, when the couplings are strong as *b*_*θ*_*/D*_*θ*_ ≫ 1, *b*_*ϕ*_*/D*_*ϕ*_ ≫ 1, and *β*_*ϕ*_*/D*_*ϕ*_ ≫ 1, we have the stable ensemble-level oscillations with *r*_*ϕ*_ ≠ 0. However, the contribution from the global coupling is proportional to 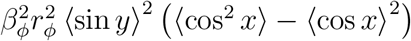, which remains minimal because ⟨sin *y*⟩^2^ ≈ 0 and ⟨cos^2^ *x*⟩−⟨cos *x*⟩^2^ ≈ 0 in the strong coupling case. Then, under the condition of *ω*_*ϕ*_ ≫ *ω*_*θ*_, components corresponding to oscillations and dragging combine to yield the following expression (Appendix F):

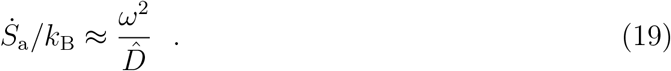

Consequently, in this strong coupling regime, the entropy production can be understood as a sum of contributions from individual oscillators, each having a renormalized oscillation frequency *ω* and noise strength 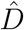. It is important to note that the contribution from the global intermolecular coupling *β*_*ϕ*_*r*_*ϕ*_ is minimal in both weak and strong coupling scenarios. This limited contribution arises because global coupling between identical molecules leads to “frictionless” interactions.

The ATPase activity, *N* ^ATP^, has been measured by counting the number of ATP molecules hydrolyzed per day per KaiC monomer [4, 5, 8]. Using the entropy production rate described in Eq. 19 with 2*π/ω*_0_ ≈ 1 day, we can write as 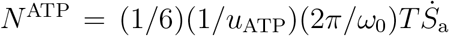, where *u*_ATP_ ~ 12*k*_B_*T* is the free energy released by the hydrolysis of one ATP molecule.

When the system is perturbed by mutations or other changes in the solution, both the oscillation frequency *ω* and the ATPase activity *N* ^ATP^ will shift from their standard values *ω*_0_ and 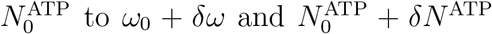, respectively. We expect that 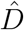 will show non-exponential variations, leading to only minor modifications. For clarity, we denote the symbol *δ* as *δ*_m_ for mutations and *δ*_T_ for temperature variations. Then, Eq. 19 leads to

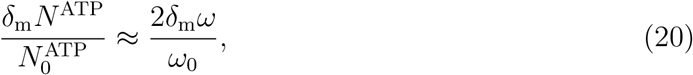

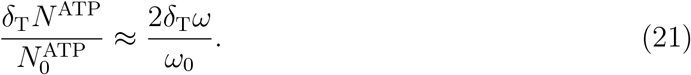

Importantly, the relationships described by Eq. 20 and 21 do not depend on model parameters and remain valid as long as the system is in the strong coupling regime. This independence from specific parameter arrangements illustrates that these correlations represent a fundamental aspect of the molecular clockwork hypothesis. Eq. 20 indicates that changes in ATPase activity, *δ*_m_*N* ^ATP^, are correlated with changes in oscillation frequency, *δ*_m_*ω*, across various mutations, as has been experimentally observed [4, 5, 8].

The ensemble-level oscillations are absent in solutions that lack KaiA or KaiB. We refer to such conditions as non-oscillatory conditions, although oscillations within each KaiC hexamer persist without synchronization in these environments in our model. These nonoscillatory conditions correspond to the scenario where *b*_*θ*_*/D*_*θ*_ ≫ 1, *b*_*ϕ*_*/D*_*ϕ*_ ≫ 1, and *β*_*ϕ*_ = 0 in the current model. Eq. 19 suggests that the impact of global coupling with *β*_*ϕ*_≠ 0 contributes only minimally to heat dissipation. Consequently, a relationship similar to Eqs. 19–21 should be observed for the ATPase activity measured under non-oscillatory conditions.

This conclusion aligns with the experimentally observed correlation between the ATPase activity in non-oscillatory conditions and the oscillation frequency across various mutations [4, 8].

When the temperature varies, the resulting ratio *δ*_T_*ω/ω*_0_ remains small, reflecting the temperature compensation for oscillations. Therefore, Eq. 21 demonstrates that ATPase activity is also temperature-compensated, as has been experimentally validated [4]. Moreover, Eq. 21 indicates that ATPase activity in non-oscillatory solutions should exhibit temperature compensation as well, a phenomenon that has also been experimentally confirmed [4, 8].

Thus, the phase model presented in this study demonstrates that the molecular clockwork hypothesis effectively accounts for the roles of CI-ATPase activity in oscillations.

### Temperature compensation, coherence, and dissipation

The present model predicts the effects of mutations and the truncation on temperature compensation, which will be important for further testing the molecular clockwork hypothesis.

For instance, the values of *Q*_10_ will change due to mutations, resulting in *Q*_10_ + *δ*_m_*Q*_10_. According to Eq. 21, the relationship 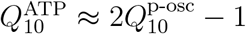 holds, where 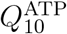 represents the *Q*_10_ for the ATPase activity, and 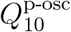 refers to the *Q*_10_ for the phosphorylation oscillations.

Therefore, Eq. 21 predicts

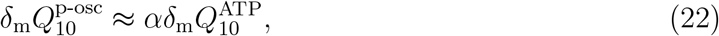

where *α* = 0.5 comes from the factor of 2 in Eq. 21. The value of *α <* 1 indicates that the temperature compensation of the oscillation frequency is less sensitive to mutations compared to the temperature compensation of ATPase activity. It is interesting to compare this coefficient with the experimentally observed value of *α* = 0.35 ± 0.13 [33], which underscores the need for further quantitative analyses of this correlation.

Another prediction of the model concerns the disconnection or truncation of the CII domain from the CI domain. The truncated CI still exhibits ATPase activity [5]. In the present model, this truncation corresponds to *b*_*θ*_ = *b*_*ϕ*_ = *ω*_*θ*_ = 0 in Eq. 18, leading to

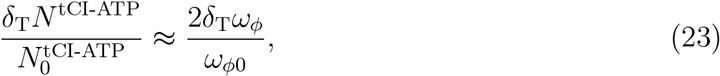

where *N* ^tCI-ATP^ represents the ATPase activity of the truncated CI domain. Since there is no frequency adjustment in *ω*_*ϕ*_ within the present model, it predicts the absence of temperature compensation in the ATPase activity of the truncated CI domain. This prediction aligns with the recently observed value of 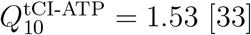.

Finally, it is important to note that heat dissipation Q in non-equilibrium processes generally comprises two components: the adiabatic component Q_a_ and the non-adiabatic component Q_na_, as Q = Q_a_+Q_na_ [40–44]. Here, Q_na_ = *T* Δ*S*_na_ represents the heat generated during the relaxation of the system from its initial state to a steadily oscillating state. We focus on the non-adiabatic entropy production per KaiC hexamer, Δ*S*_na_, arising from the change in the distribution from *P* ({*θ, ϕ*}) to the steadily oscillating state *P*_0_({*θ, ϕ*}) of the standard solution, which is expressed as Δ*S*_na_*/k*_B_ = (1*/N*_c_) *dθdϕP* ({*θ, ϕ*}) ln [*P* ({*θ, ϕ*})*/P*_0_({*θ, ϕ*})] [44]. Using the mean-field approximation and starting from a non-oscillatory state with a small *β*_*ϕ*_, the entropy required to achieve a steadily oscillating state with a larger *β*_*ϕ*_ is

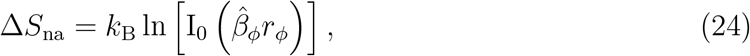

where I_0_(· · ·) is the modified Bessel function of the first kind, and 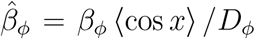 is the effective coupling constant at the steadily oscillating state (Appendix G). Q_na_ = *T* Δ*S*_na_ is the free-energy cost required to transition the system from a non-oscillatory state to a steadily oscillating state. This free-energy cost is necessary to ensure the system’s robustness against perturbations, such as the temporary inactivation of KaiA or KaiB.

In the steadily oscillating state, the effective coupling 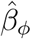 is large and *r*_*ϕ*_ ≈ 1. Therefore, Eq. 24 can be expressed as 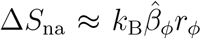. By combining this expression with Eq. 9 and using 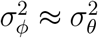, we arrive at the relation:

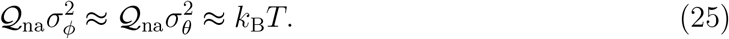

This relation indicates a trade-off between the free-energy cost of robustness and the coherence of the ensemble-level oscillations that are achieved. Importantly, this prediction is independent from the precise selection of parameters, indicating that this trade-off is based on a fundamental feature of the molecular clockwork hypothesis. Therefore, Eq. 25 could provide a basis for further testing the molecular clockwork hypothesis.

## DISCUSSION

Biochemical and structural analyses have highlighted the importance of CI-ATPase reactions [4–8], leading to the hypothesis that these reactions drive the pacemaker mechanism within the KaiC hexamer. This hypothesis was illustrated by Kondo using a metaphor from artificial clockwork [2, 3], which sparked significant interest and emphasized the need for further quantitative assessment.

In this study, we introduced a theoretical model to evaluate this molecular clockwork hypothesis and demonstrated its effectiveness. It is important to note that our approach differs in several ways from Kondo’s original proposal of the hypothesis [2, 3]. Kondo described the ATP-driven structural oscillator in KaiC as a “harmonic” oscillator, likening it to a “pendulum.” In contrast, we characterize this oscillator as a limit cycle generated by a nonlinear mechanism within the feedback relationships between the CI-ATPase reactions and the structural transitions. Additionally, Kondo referred to the phosphorylation oscillations as a “mainspring” that supplies energy to the oscillations occurring within each molecule, since the phosphorylation oscillations should largely stabilize the pacemaker oscillations. Also in our model, the phosphorylation oscillations in *θ* stabilize the pacemaker oscillations in *ϕ* through the intermolecular coupling *β*_*ϕ*_*r*_*ϕ*_ sin(*ϕ* − ⟨*ϕ*⟩) cos(*θ* − *ϕ*). However, this stabilization does not directly contribute to the free-energy balance at the steadily oscillating state, and we propose that ATP consumption in CI serves as the free-energy source for these oscillations.

With the current approach, there are still many intriguing issues that require further investigation. First, the phase model should be utilized to determine if it can explain the phase entrainment observed when two oscillating KaiABC solutions with different phases are mixed [46]. Additionally, it is crucial to examine the phase response to sudden changes in ATP or ADP concentration [47], or to changes in temperature [48].

Furthermore, it is particularly interesting to extend the model to analyze the fluctuations among monomer subunits within the KaiC hexamer [49]. We denote the phase variable representing the structural state of the *k*th subunit in the *i*th KaiC hexamer as *ϕ*_*ik*_ with *k* = 1 ~ 6. Then, we can express the relationships between *ϕ*_*ik*_ and the overall structural phase *ϕ*_*i*_ of the hexamer as

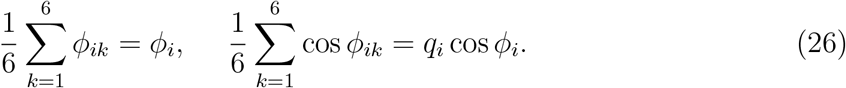

In this study, we have assumed that the distribution of *ϕ*_*ik*_ is narrowly peaked around *ϕ*_*i*_, resulting in *q*_*i*_ = 1; however, it is generally more accurate to consider that *q*_*i*_ *<* 1. In this scenario, the pacemaker frequency of each subunit, 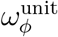, must be renormalized by coupling constants that represent the interactions between subunits, to give rise to the pacemaker frequency of the hexamer, *ω*_*ϕ*_, which was the main focus of our current research. This framework should help clarify the diverse effects of mutations that alter subunit-subunit interactions within the KaiC hexamer, which can result in various modifications to the intramolecular couplings in the clockwork mechanism. Furthermore, this framework should provide a more quantitative explanation for temperature compensation.

Additionally, it is crucial to derive the values of coupling strengths from the atomistic point of view. In the present study, the intramolecular interactions between the structural state and the phosphorylation level, as well as the global intermolecular interaction, were defined phenomenologically using parameters *b*_*θ*_, *b*_*ϕ*_, and *β*_*ϕ*_. Deriving these parameters from detailed atomistic knowledge through molecular simulations or spectroscopy is an important direction for further research. Such microscopic derivations should enable predictions regarding the effects of additional mutations in KaiC.

## CONCLUSION

We have developed a theoretical model to assess the effectiveness of the molecular clock-work hypothesis. Our findings illustrate the hypothesis’s ability to explain several key observations. The mean-field structure of our model shows how the frequency of intramolecular structural oscillations is renormalized by molecular coupling, thereby accounting for the observed correlation between oscillation frequency and ATPase activity across various mutations in the KaiC protein. Importantly, we addressed the significant changes (over 10-fold) in the oscillation period that result from mutations located at the domain boundary in KaiC. Furthermore, the mean-field renormalization scheme clarifies the mechanisms that underlie temperature compensation in both the oscillation period and ATPase activity of KaiC. Our model also predicts the quantitative effects of mutations and truncations on temperature compensation and suggests a relationship between heat dissipation and oscillation coherence, paving the way for further tests of our hypothesis.

We based our model on two fundamental assumptions: (1) The structural oscillations driven by the CI-ATPase reactions serve as the primary pacemaker for oscillations, with *ω*_*ϕ*_ ≫ *ω*_*θ*_. (2) The pacemaker couplings, *b*_*θ*_ and *b*_*ϕ*_, work together, with *b*_*θ*_ being more sensitive to temperature and the side-chain volume of the residue at the CI-CII boundary than *b*_*ϕ*_. Although these fundamental assumptions have not yet been experimentally validated directly, the theoretical predictions stemming from them are consistent with many observations, providing clear explanations of the mechanisms involved. These results should motivate further comparisons with experimental analyses, which can be achieved particularly through molecular dynamics simulations and structural measurements focused on the escapement coupling mechanisms.

Thus, investigating the molecular clockwork hypothesis will deepen our understanding of how ensemble-level macro-or mesoscopic oscillations emerge and are regulated by the microscopic atomic structure of proteins.

## APPENDIX A

**PACEMAKER OSCILLATIONS INDUCED BY REACTIONS AND STRUCTURAL TRANSITIONS**

We present a simple model to explain a potential mechanism by which the coupled dynamics of CI-ATPase reactions and structural transitions can induce oscillations. We define the probability that the *k*th subunit of a KaiC hexamer adopts the *p* structure as 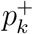 with *k* = 0, …, 5, and the probability of the subunit adopting the *dp* structure as 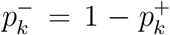. Additionally, we express the probability that CI of the *k*th subunit binds ATP as 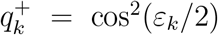, and the probability that CI of the *k*th subunit binds ADP as 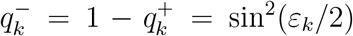. Therefore, 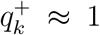 when *ε*_*k*_ ≈ 2*nπ*, and 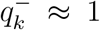 when *ε*_*k*_ ≈ (2*n* + 1)*π*, where *n* is an integer.

We assume that the *p* structure is stabilized when ATP binds to CI, while the *dp* structure is stabilized when ADP binds to CI. In this framework, the release of Pi from CI is expected to trigger the transition from the *p* structure to the *dp* structure, which should occur at *ε*_*k*_ ≈ (2*n*+1*/*2)*π* with sin *ε*_*k*_ ≈ 1. Similarly, the release of ADP, followed by the incorporation of ATP, should facilitate the transition from the *dp* structure back to the *p* structure, which is expected to happen at *ε*_*k*_ ≈ (2*n* + 3*/*2)*π* with sin *ε*_*k*_ ≈ −1. Furthermore, we assume that the neighboring subunits in the KaiC hexamer tend to take the same structure. We write the neighboring subuits of the *k*th subunit, *k*_*p*_ and *k*_*m*_ with *k*_*p*_ = mod(*k* + 1, 6) and *k* = mod(*k*_*m*_ + 1, 6).

Then, using constants, *σ* > 0, *ρ* > 0, *ζ* > 0, and *η* > 0, we describe the stochastic process governing these structural transitions as

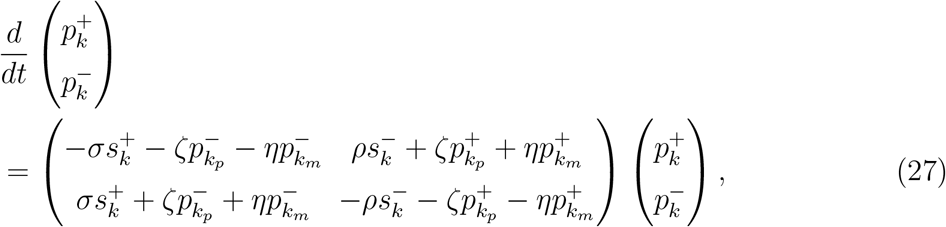

where 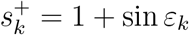 and 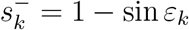

Additionally, as nucleotide binds near the interface between the *k*th and *k*_*p*_th subunits, we assume that the Pi release, which marks the transition from 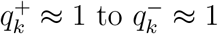, is promoted when the *k*th and *k*_*p*_th subunits adopt the *p* structure with 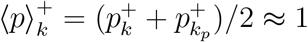, while the ADP release and the following ATP incorporation, which is the transition from 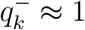 to 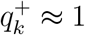, is promoted when CI adopts the *dp* structure with 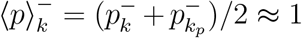. Similar to modeling mechano-chemical coupling in molecular motors, we represent this scenario with the following Langevin equation:

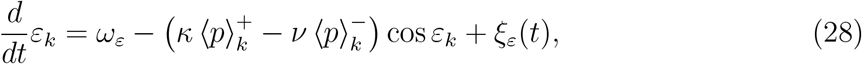

where *κ* > 0 and *ν* > 0 are constants represnting the feedback effect of the CI structure on the reactions, and *ξ*_*ε*_ represnts the noise effect, satisfying ⟨*ξ*_*ε*_(*t*)⟩ = 0 and ⟨*ξ*_*ε*_(*t*)*ξ*_*ε*_(*t*^*′*^)⟩ = 2*D*_*ε*_*δ*(*t* − *t*^*′*^), with a constant *D*_*ε*_ > 0. *ω*_*ε*_ > 0 is the driving force of ATPase reactions with log(*ω*_*ε*_*/D*_*ε*_) ∝ (*µ*_ATP_ − (*µ*_ADP_ + *µ*_Pi_)) */*(*k*_B_*T*), where *k*_B_ is the Boltzmann constant, *T* is temperature, and *µ*_ATP_, *µ*_ADP_, and *µ*_Pi_ are chemical potentials of ATP, ADP, and Pi, respectively. When *ω*_*ε*_ is sufficiently large, trapping of the reactions to *ε*_*k*_ ≈ (*n* + 1*/*2)*π* is avoided, allowing cycles of reactions to be maintained. This persistence, in turn, drives the structural oscillations through the coupling shown in Eqs. 27 and 28. An example trajectory representing these oscillations is illustrated in Fig. 5. In this trajectory, the probability of ADP release (the probability to show 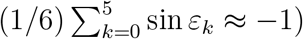 is large during the phosphorylation phase in the *p* structure of 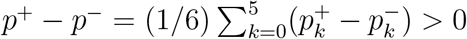, which aligns with the experimental observations [4]. Further detailed simulations of the coupled dynamics of reactions and structural transitions in the KaiC hexamer were discussed in previous reports [9, 10, 22, 23].

**FIG. 5.**
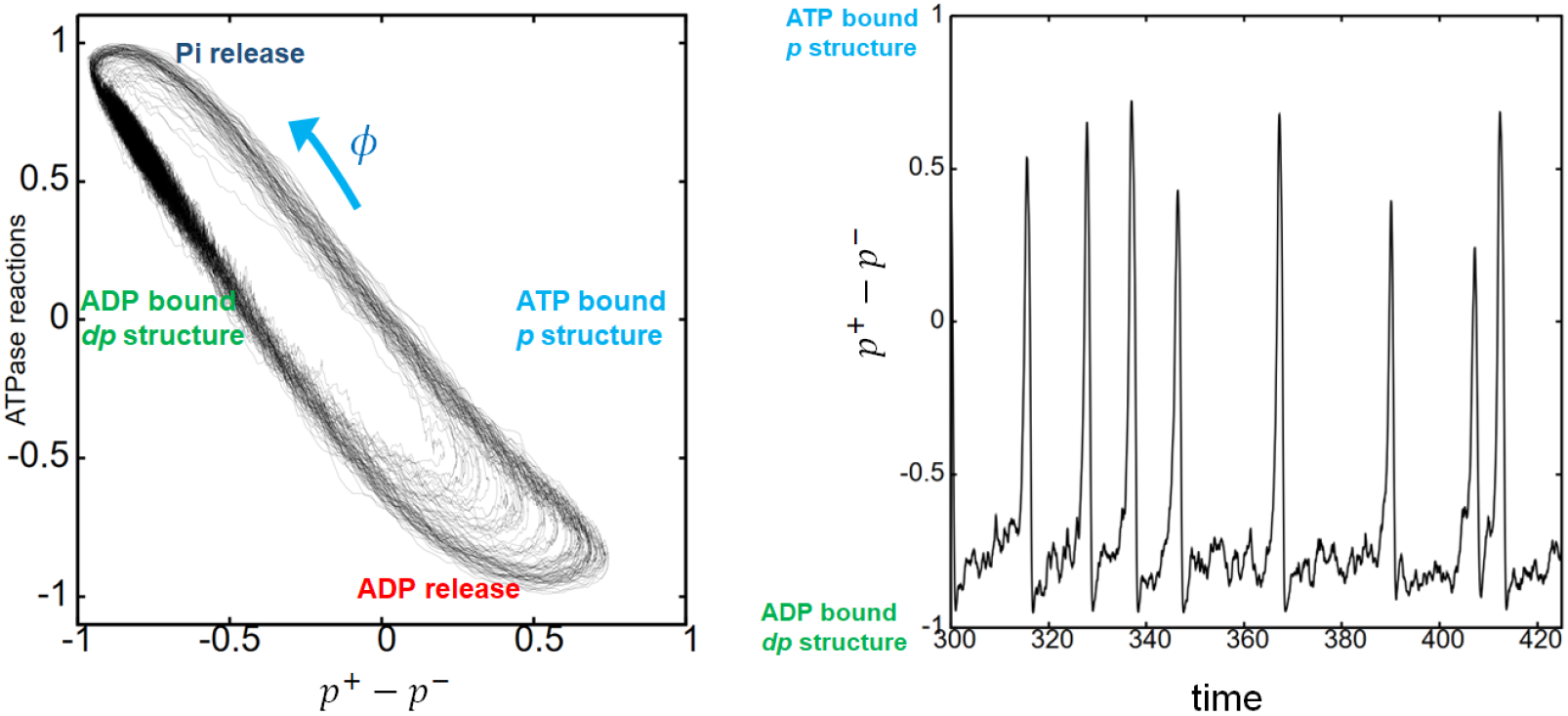
Pacemaker oscillations induced by the coupled dynamics of ATPase reactions and structural transitions. (Left) An example trajectory of the simulated oscillations is plotted on the plane of the structural state, *p*^+^ *p*^−^, and the ATPase reaction state, 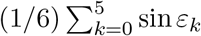 where *p*^+^ is the probability that the KaiC hexamer shows the *p* structure, 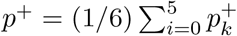, *p*^+^, and *p*^−^ = 1 − *p*^+^. The *p* structure predominantly appeas when *p*^+^ −*p*^−^ > 0, while the *dp* structure becomes dominant when *p*^+^ − *p*^−^ *<* 0. The phase of this limit cycle, *ϕ*, serves as the coordinate for the pacemaker oscillations discussed in the main text. (Right) The structural oscillations depicted in the left panel are shown as a function of time. Parameters used are: *σ* = 4, *ρ* = 2, *ζ* = 20, *η* = 10, *κ* = *ν* = 5, *D*_*ε*_ = 0.1, and *ω*_*ε*_ = 3.

In the circular trajectory illustrated in Fig. 5, we can define the phase *ϕ* along this trajectory [32]. When we write the *ϕ*-dependent driving force of these oscillations as 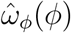, the equation of motion of the phase should become

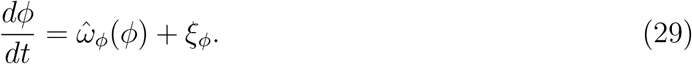

When we approximate the frequency as a constant by setting 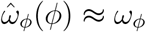, Eq. 29 simplifies to the phase equation presented in Eq. 2 with *b*_*ϕ*_ = *F* (*θ, ϕ*) = 0. In this manner, the current Appendix provides an example mechanism that can account for the origin of the pacemaker oscillations discussed in the main text. However, the assumptions utilized in Eqs. 27 and 28 should be examined more closely, and the further research should be needed for constructing the suitable model for the origin of the pacemaker oscillations.

## APPENDIX B

**CONCENTRATION OF FREE, UNBOUND KAI A DIMERS**

We write total concentration of the KaiC hexamers as *c*. The concentration of the KaiC hexamers having the *dp* structure is *c*^*dp*^ = *c* (1 − *r*_*ϕ*_ cos ⟨*ϕ*⟩) */*2. We write the concentration of the *dp*-structure hexamer that does not bind a KaiA dimer on the CI-bound KaiB as 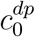 and the concentration of the *dp*-structure hexamer that does bind as 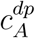, so that 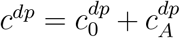. The total concentration of the KaiA dimers is *a*, and the concentration of free KaiA dimers that are unbound from KaiC or KaiB is *a*_0_. We approximate that each KaiA binds to each KaiB monomer independently. Then, using the dissociation constant, 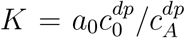, and considering that six KaiB monomers bind to the CI of the KaiC hexamer, we have

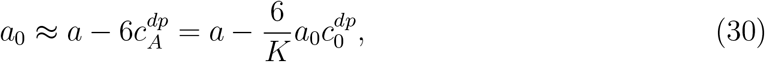

which represents the strong tendency of KaiA sequestration to the KaiA-KaiB-KaiC complex. We also have

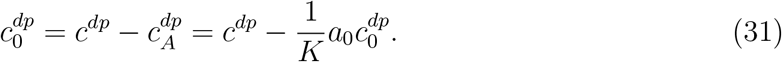

Eqs. 30 and 31 lead to 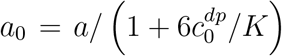 and 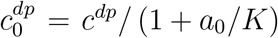; combining these equations leads to the expression Eq. 3.

In Fig.6, we plot examples of oscillations in the concentration of free KaiA dimers *a*_0_ as a function of the phase of the collective ensemble-level oscillations ⟨*ϕ*⟩. These oscillations are well fitted by the function *a*_0_ ≈ *γ*_0_ + *γ*_1_*r*_*ϕ*_ cos⟨*ϕ*⟩. Fig.6 shows that *γ*_1_*r*_*ϕ*_ ≪ *c* for a small *a* and *γ*_0_ ≫ *γ*_1_*r*_*ϕ*_ for a large *a*, showing that the *a*_0_ oscillations are effective only in the limited range of the total concentration of KaiA dimers *a*.

**FIG. 6.**
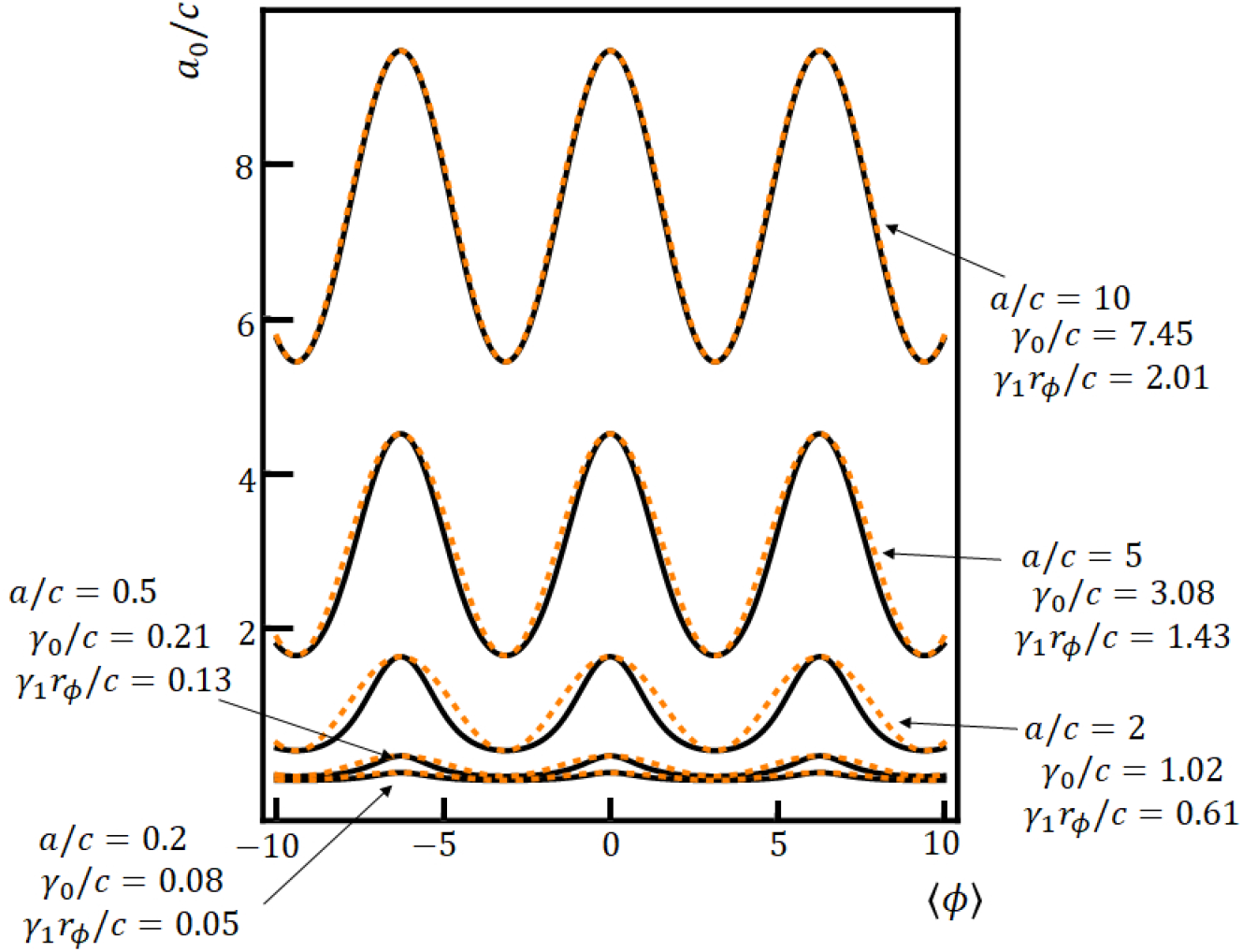
Examples of oscillations in the concentration of free KaiA dimers *a*_0_, normalized by the total concentration of KaiC hexamers *c*, are shown as a function of the phase of collective oscillations of the ensemble of KaiC hexamers ⟨*ϕ*⟩, for various total concentrations of KaiA dimers *a*, respectively. The black lines represent *a*_0_*/c*, which were derived from Eq.3 for the values *a/c* = 0.2, 0.5, 2.0, 5.0, and 10. The dissociation constant was set to be *K/c* = 1 and a fixed value of the order parameter, *r*_*ϕ*_ = 0.8, was assumed. Here, *a*_0_ can be well approximated by *a*_0_ ≈ *γ*_0_+*γ*_1_*r*_*ϕ*_ cos⟨*ϕ*⟩ (orange dashed lines), showing that *γ*_1_*r*_*ϕ*_ is small when *a/c* is low, while *γ*_0_*/*(*γ*_1_*r*_*ϕ*_) grows large when *a/c* is high.

## APPENDIX C

**MEAN-FIELD APPROXIMATION**

Using the mean-field factorization, 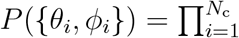 *P*_*x*_(*x*_*i*_)*P*_*y*_(*u*_*i*_), with *x*_*i*_ = *θ*_*i*_−*ϕ*_*i*_ and *y*_*i*_ = *ϕ*_*i*_−⟨*ϕ*⟩, the Fokker-Planck equation (Eq. 7) becomes

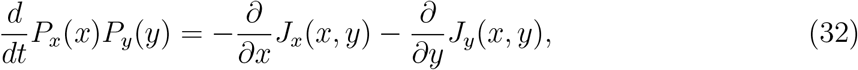

with

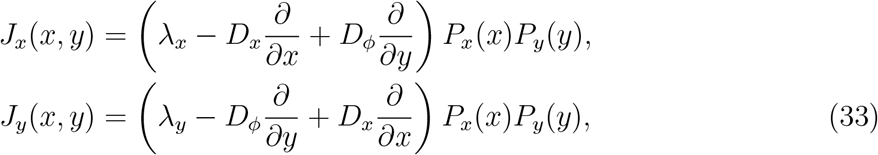

where *D*_*x*_ = *D*_*θ*_ + *D*_*ϕ*_ and

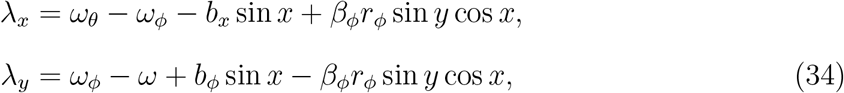

with *b*_*x*_ = *b*_*θ*_ +*b*_*ϕ*_. Eq. 32 describes fluctuations *x* and *y* around the ensemble-level oscillations ⟨*ϕ*⟩ = *ωt*. For realizing the steadily oscillating states, *P*_*x*_(*x*) and *P*_*y*_(*y*) need to have static solutions with limited distribution widths.

By integrating Eq. 32 with the variable *x* and using the relation *ω*_*ϕ*_ − *ω* + *b*_*ϕ*_ ⟨sin *x*⟩ = 0, we obtain

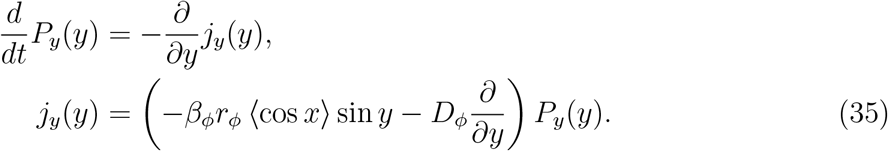

Eq. 35 has the static periodic solution,

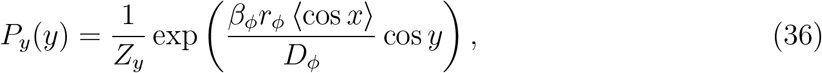

with

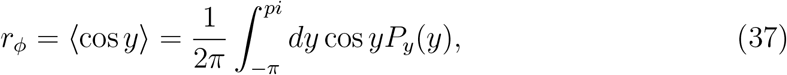

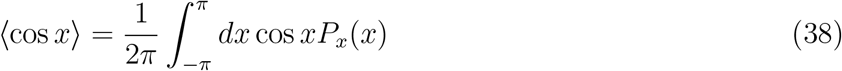

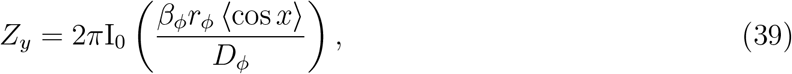

where I_0_(· · ·) in Eq. 39 is the modified Bessel function of the first kind. Eq. 37 represents the self-consistent relation for *r*_*ϕ*_ and *P*_*y*_(*y*). With Eq. 36, we have ⟨*y*⟩ = ⟨sin *y*⟩ = 0, and the probability current vanishes to be *j*_*y*_ = 0.

By integrating Eq. 32 with the variable *y* and using ⟨sin *y*⟩ = 0, we obtain

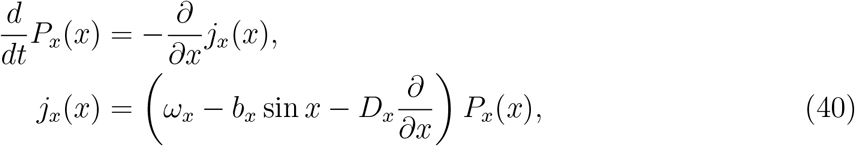

where *ω*_*x*_ = *ω*_*θ*_ − *ω*_*ϕ*_. Eq. 40 has the static solution having the periodicity of *P*_*x*_(*x* + 2*π*) = *P*_*x*_(*x*) [50] as

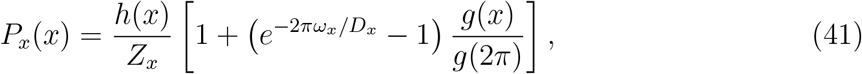

where

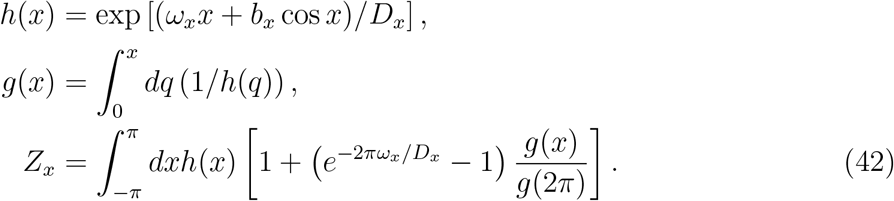

It is important to note that even with this static solution, the probability current of Eq. 40 does not vanish, showing 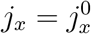 with

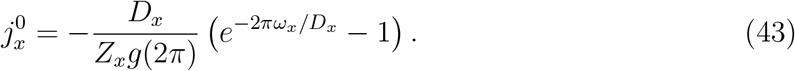

In other words, even when the system reaches a steadily oscillating state, as described by Eq. 41, the fluctuation *x* = *θ* − *ϕ* exhibits a biased stochastic motion. Due to the periodic nature of the system with respect to *x*, the distribution *P*_*x*_(*x*) remains static because of the balance between incoming and outgoing fluxes of the constant current, 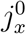. We have ⟨*x*⟩≠ 0 and ⟨sin *x*⟩≠ 0. *P*_*x*_ becomes narrower as the ratio *b*_*x*_*/D*_*x*_ increases.

In Fig. 2, the mean-field results were obtained by numerically solving the self-consistent relation Eq. 37, from which *P*_*y*_(*y*) in Eq. 36 was derived. ⟨cos *x*⟩ was numerically calculated in Eq. 38. Other expectation values were numerically calculated by using the thus obtained *P*_*y*_(*y*) and *P*_*x*_(*x*).

Eqs. 35 and 40 have the same form,

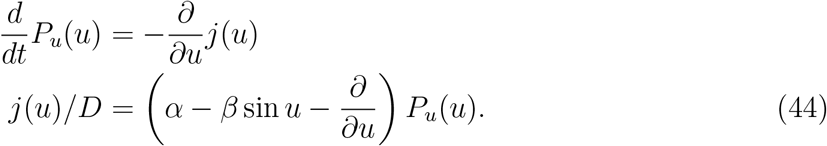

Eq. 40 for *u* = *x* is recovered by putting *α* = *ω*_*x*_*/D*_*x*_, *β* = *b*_*x*_*/D*_*x*_, and *D* = *D*_*x*_, and Eq. 35 for *u* = *y* is recovered by putting *α* = 0, 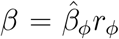, and *D* = *D*_*ϕ*_ into Eq. 44. Eq. 44 has a static solution [50],

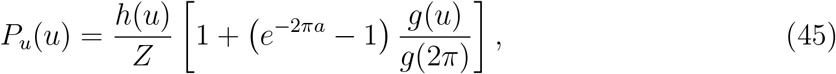

where *h*(*u*) = exp (*αx* + *β* cos *x*). Using Eq. 45, expectation values are calculated as 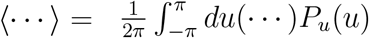Some of them can be expressed explicitly in the weak coupling (*β* ≪ 1) and in the strong coupling (*β* ≫ 1) cases as

- **Weak coupling case (***β* ≪ 1)

In this case, we have the following expressions:

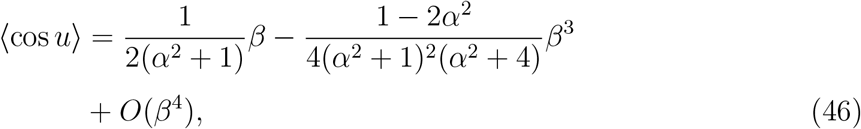

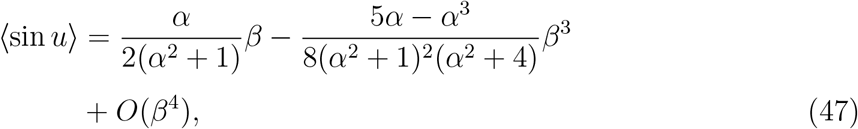

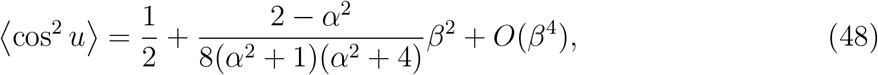

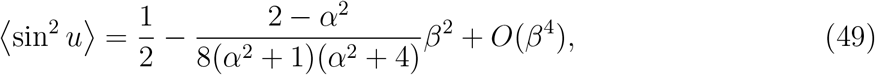

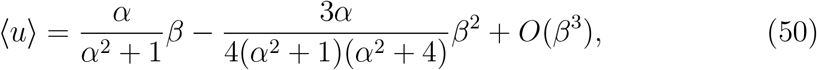

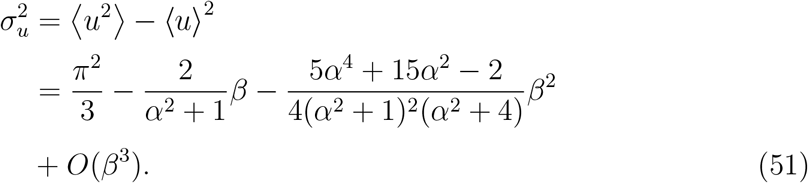

1. **Strong coupling case (***β* ≫ 1)

In this case, *P*_*u*_(*u*) can be approximated by a Gaussian,

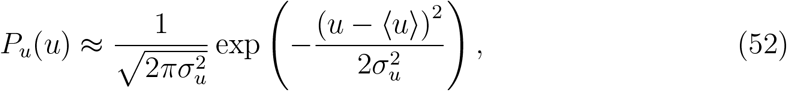

where

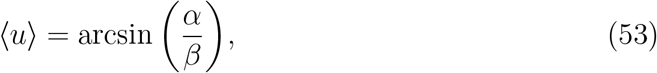

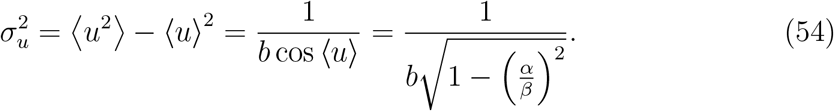

Using this expression for *P*_*u*_(*u*), we have

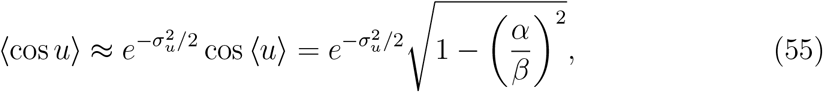

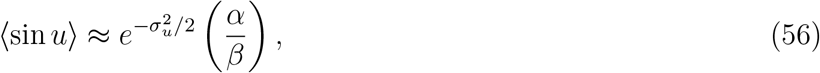

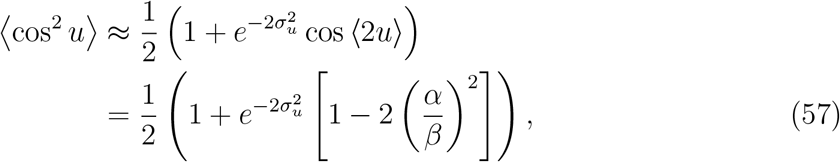

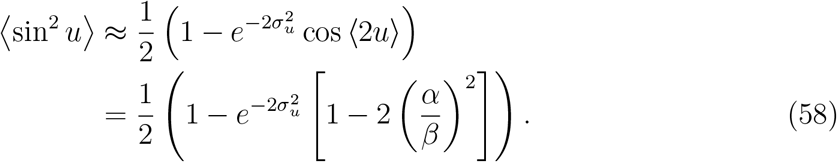

## APPENDIX D

**TRANSITION FROM THE NON-OSCILLATORY TO OSCILLATORY STATES**

The global ensemble-level oscillations are realized when *r*_*ϕ*_ ≠ 0. Here, the order parameter *r*_*ϕ*_ is defined by ⟨cos *ϕ*⟩ = *r*_*ϕ*_ cos ⟨*ϕ*⟩. Using *ϕ* = *y* + *ωt*, we have ⟨cos(*y* + *ωt*)⟩ = ⟨cos *y*⟩ cos *ωt* − ⟨sin *y*⟩ sin *ωt* = *r*_*ϕ*_ cos ⟨*y*⟩ cos *ωt* − *r*_*ϕ*_ sin ⟨*y*⟩ sin *ωt*. Because ⟨sin *y*⟩ = 0 and ⟨*y*⟩ = 0, this definition becomes *r*_*ϕ*_ = ⟨cos *y*⟩. By putting *α* = 0 and 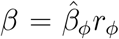 into Eq. 46 for *u* = *y*, we have

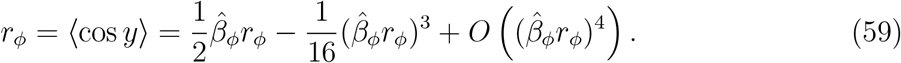

This is the self-consis tent rela tion to determine *r*_*ϕ*_. For 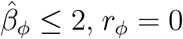 is the unique solution. When we neglect 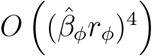 for 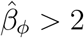, we have another solution,

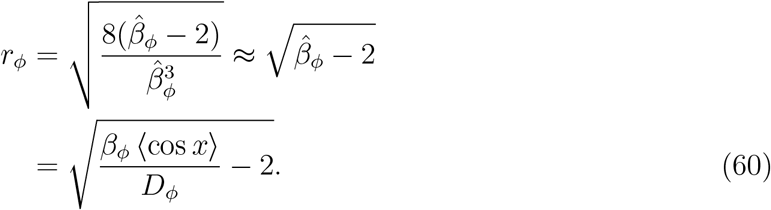

Therefore, the increase in the coupling strength from 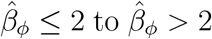 is the transition from the non-oscillatory to oscillatory states, where *r*_*ϕ*_ shows a critical behavior as in Eq. 60. For the larger coupling strength 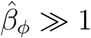, from Eq. 55, we find *r*_*ϕ*_ approaches to 1 as

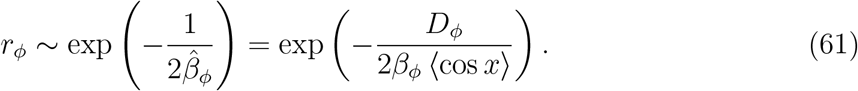

## APPENDIX E

**VARIANCES AMONG OSCILLATORS**

When *r*_*ϕ*_ = 0, we have *P*_*y*_(*y*) = 1*/*(2*π*), which corresponds to the *β* = 0 case in Eq. 51, resulting in 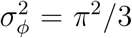. When *r*_*ϕ*_ is positive, by putting *α* = 0 and 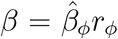 in Eq. 51, we find 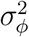 is reduced to 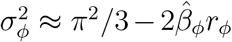. When the coupling 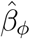 is large enough, from Eq. 54, we have 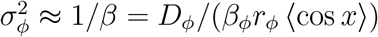.

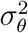 is calculated by

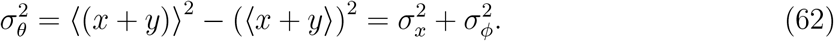

when the couplings, *b*_*θ*_ and *b*_*ϕ*_, are large, we find 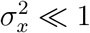 and 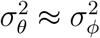.

## APPENDIX F

**ENTROPY PRODUCTION RATE AT THE STEADILY OSCILLATING STATE**

In the mean-field expression, currents defined in Eq. 8 in the main text are

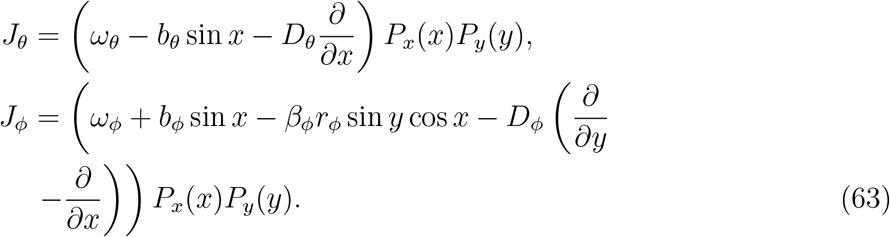

Using Eq. 63, the house-keeping [40] or the adiabatic [42–44] component of the entropy production rate at the steadily oscillating state is

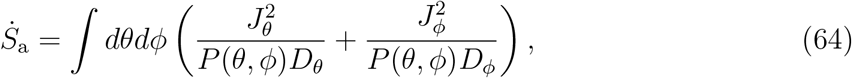

By putting *P* (*θ, ϕ*) = *P*_*x*_(*x*)*P*_*y*_(*y*) into this relation, we have the following expression:

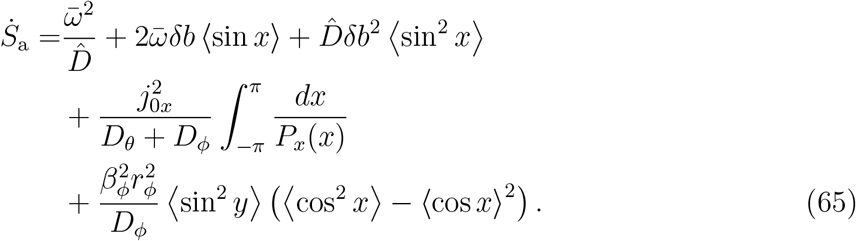

Here,

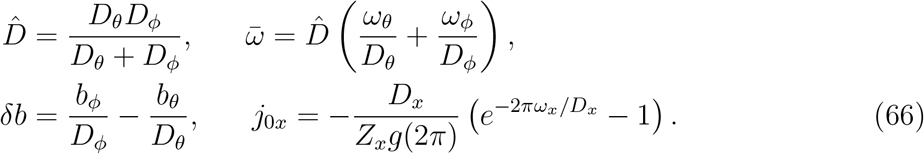

The term proportional to 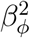 in Eq. 65 becomes zero when *r*_*ϕ*_ = 0. This situation occurs either when the coupling *β*_*ϕ*_ is small, resulting in a loss of global synchronization, or when the couplings *b*_*θ*_ and *b*_*ϕ*_ are small, which leads to a small value of ⟨cos *x*⟩. Conversely, when *β*_*ϕ*_, *b*_*θ*_, and *b*_*ϕ*_ are large, we anticipate *r*_*ϕ*_ ≈ 1 and *r*_*x*_ = ⟨cos *x*⟩ */* cos ⟨*x*⟩ ≈ 1. However, from Eqs. 55, 57, and 58, The term proportinal to 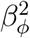 becomes

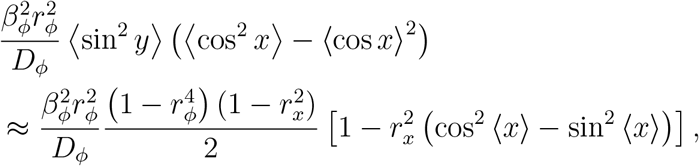

which also vanishes when *r*_*ϕ*_ ≈ 1 and *r*_*x*_ ≈ 1. The other terms in Eq. 65 are calculated using the expressions for *P*_*x*_(*x*), ⟨sin *x*⟩, and sin *x*. To represent them explicitly, we can utilize Eqs. 47 and 49 in the weak coupling case, as well as Eqs. 56 and 58 in the strong coupling case.

In the weak coupling case, the term proportional to 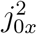 in Eq. 65 becomes

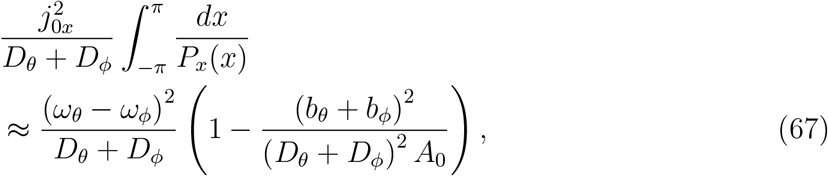

where

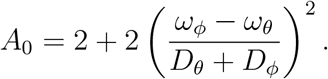

Using Eqs. 47 and 49 in the first line of Eq. 65, retaining the terms up to 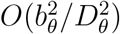 and 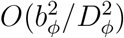, and combining it with the expression of Eq. 67, we have

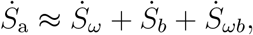

where

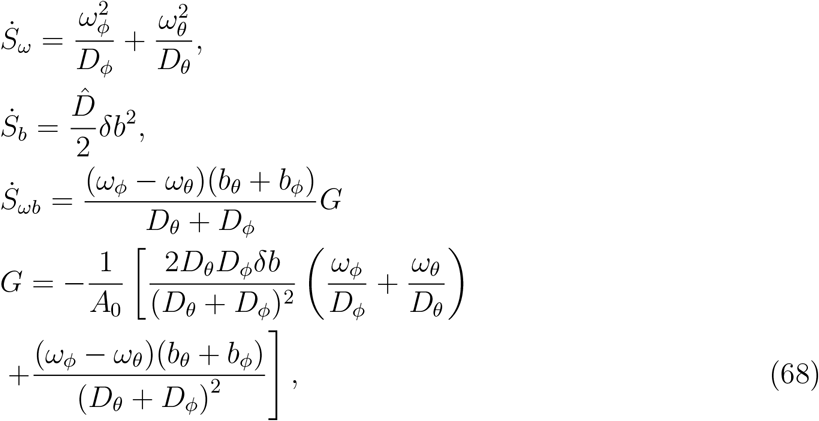

which leads to Eq. 14 in the main text.

In the strong coupling case, the term proportional to 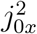 in Eq. 65 vanishes because *j*_0*x*_ ≈ 0. Then, using Eqs. 56 and 58 in the first line of Eq. 65, we have

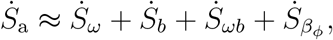

where

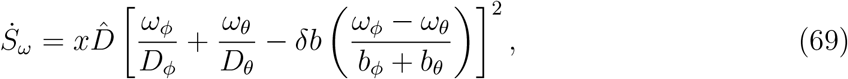

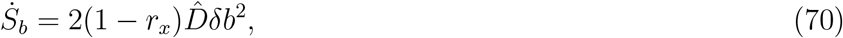

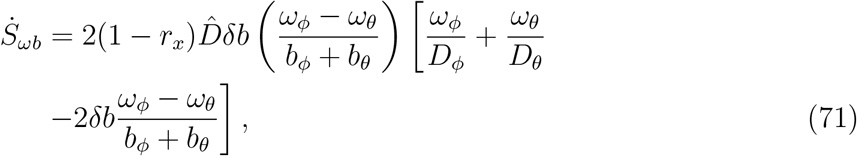

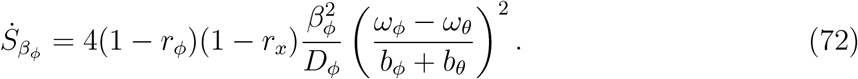

Then, in the strong coupling limit, neglecting the terms proportional to 1 − *r*_*ϕ*_ or 1 − *r*_*x*_, we have 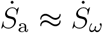. When the activity in the structural transitions driven by the CI-ATPase reactions is large with *ω*_*ϕ*_ ≫ *ω*_*θ*_, this expression leads to

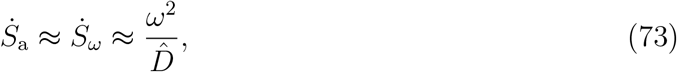

which is Eq. 15 in the main text.

## APPENDIX G

**NON-ADIABATIC ENTROPY PRODUCTION**

We consider the scenario that due to a sudden decrease in *β*_*ϕ*_, the system reaches a non-oscillatory state with *r*_*ϕ*_ = 0. Then, the system returns to the oscillatory state when *β*_*ϕ*_ recovers to have a large value that allows for *r*_*ϕ*_ > 1. We calculate the non-adiabatic entropy required for this recovery process. During this process, we assume that the intramolecular couplings, *b*_*θ*_ and *b*_*ϕ*_, remain large to suppress the *x* fluctuation. In the non-oscillatory state with *r*_*ϕ*_ = 0, we have 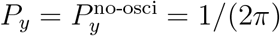. Then, using the mean-field approximation and *Z*_*y*_ of Eq. 39, the non-adiabatic entropy production per KaiC hexamer is

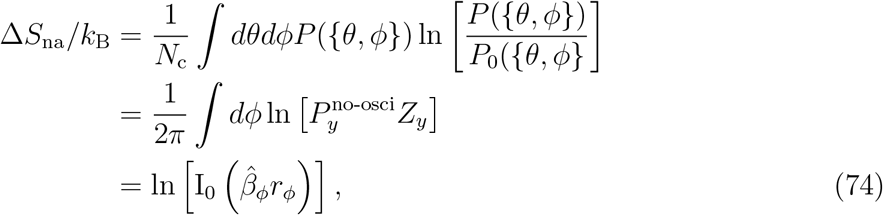

where 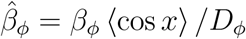. When 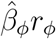 is large, we have 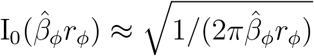 exp 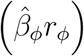. In this scenario, by dropping the term proportional to ln 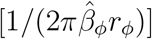, we have 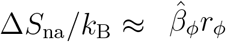, which leads to Eq. 25.

## ACKNOWLEDGEMENT

The authors would like to thank Prof. Shuji Akiyama for discussions highlighting the effects of mutations and truncation on temperature compensation. This work was supported by JSPS-KAKENHI Grants 22H00406, 24H00061, and 25K24664.

## References

[1] M. Nakajima, K. Imai, H. Ito, T. Nishiwaki, Y. Murayama, H. Iwasaki, T. Oyama, and T. Kondo, Reconstitution of circadian oscillation of cyanobacterial KaiC phosphorylation in vitro, Science 308, 414 (2005).

[2] T. Kondo, Around the circadian clock: Review and preview, in Circadian Rrhythms in Bacteria and Microbiomes, edited by C. H. Johnson and M. J. Rust (Springer, 2021) pp. 21–52.

[3] K. Ito-Miwa, K. Terauchi, and T. Kondo, Mechanism of the cyanobacterial circadian clock protein KaiC to measure 24 hours, in Circadian Rrhythms in Bacteria and Microbiomes, edited by C. H. Johnson and M. J. Rust (Springer, 2021) pp. 123–130.

[4] K. Terauchi, Y. Kitayama, T. Nishiwaki, K. Miwa, Y. Murayama, T. Oyama, and T. Kondo, ATPase activity of KaiC determines the basic timing for circadian clock of cyanobacteria, Proc. Natl. Acad. Sci. U S A 104, 16377 (2007).

[5] J. Abe, T. B. Hiyama, A. Mukaiyama, S. Son, T. Mori, S. Saito, M. Osako, J. Wolanin, E. Yamashita, T. Kondo, and S. Akiyama, Atomic-scale origins of slowness in the cyanobacterial circadian clock, Science 349, 312 (2015).

[6] C. Phong, J. S. Markson, C. M. Wilhoite, and M. J. Rust, Robust and tunable circadian rhythms from differentially sensitive catalytic domains, Proc. Natl. Acad. Sci. U S A 110, 1124 (2013).

[7] Y. Furuike, A. Mukaiyama, S. I. Koda, D. Simon, D. Ouyang, K. Ito-Miwa, S. Saito, E. Yamashita, T. Nishiwaki-Ohkawa, K. Terauchi, T. Kondo, and S. Akiyama, Regulation mechanisms of the dual ATPase in KaiC, Proc. Natl. Acad. Sci. U S A 119, e2119627119 (2022).

[8] K. Ito-Miwa, Y. Furuike, S. Akiyama, and T. Kondo, Tuning the circadian period of cyanobacteria up to 6.6 days by the single amino acid substitutions in KaiC, Proc. Natl. Acad. Sci. U S A 117, 20926 (2020).

[9] J. Paijmans, D. K. Lubensky, and P. R. Ten Wolde, A thermodynamically consistent model of the post-translational Kai circadian clock, PLoS Comput. Biol. 13, e1005415 (2017).

[10] M. Sasai, Role of the reaction-structure coupling in temperature compensation of the KaiABC circadian rhythm, PLoS Comput. Biol. 18, e1010494 (2022).

[11] E. Emberly and N. S. Wingreen, Hourglass model for a protein-based circadian oscillator, Phys. Rev. Lett. 96, 038303 (2006).

[12] T. Mori, D. R. Williams, M. O. Byrne, X. Qin, M. Egli, H. S. McHaourab, P. L. Stewart, and C. H. Johnson, Elucidating the ticking of an in vitro circadian clockwork, PLoS Biol. 5, e93 (2007).

[13] M. Yoda, K. Eguchi, T. P. Terada, and M. Sasai, Monomer-shuffling and allosteric transition in KaiC circadian oscillation, PLoS One 2, e408 (2007).

[14] K. Eguchi, M. Yoda, T. P. Terada, and M. Sasai, Mechanism of robust circadian oscillation of KaiC phosphorylation in vitro, Biophys. J. 95, 1773 (2008).

[15] D. Zhang, Y. Cao, Q. Ouyang, and Y. Tu, The energy cost and optimal design for synchronization of coupled molecular oscillators, Nat. Phys. 16, 95 (2020).

[16] H. Kageyama, T. Nishiwaki, M. Nakajima, H. Iwasaki, T. Oyama, and T. Kondo, Cyanobacterial circadian pacemaker: Kai protein complex dynamics in the KaiC phosphorylation cycle in vitro, Mol. Cell 23, 161 (2006).

[17] H. Takigawa-Imamura and A. Mochizuki, Predicting regulation of the phosphorylation cycle of KaiC clock protein using mathematical analysis, J. Biol. Rhythms 21, 405 (2006).

[18] J. S. van Zon, D. K. Lubensky, P. R. Altena, and P. R. ten Wolde, An allosteric model of circadian KaiC phosphorylation, Proc. Natl. Acad. Sci. U S A 104, 7420 (2007).

[19] T. S. Hatakeyama and K. Kaneko, Generic temperature compensation of biological clocks by autonomous regulation of catalyst concentration, Proc. Natl. Acad. Sci. U S A 109, 8109 (2012).

[20] M. J. Rust, J. S. Markson, W. S. Lane, D. S. Fisher, and E. K. O’Shea, Ordered phosphorylation governs oscillation of a three-protein circadian clock, Science 318, 809 (2007).

[21] Y. K. Lee and C. Hyeon, Physical constraints on the rhythmicity of the biological clock, Phys. Rev. Res. 10.1103/9byx-f411 (2026).

[22] S. Das, T. P. Terada, and M. Sasai, Role of ATP hydrolysis in cyanobacterial circadian oscillator, Sci. Rep. 7, 17469 (2017).

[23] M. Sasai, Mechanism of autonomous synchronization of the circadian KaiABC rhythm, Sci. Rep. 11, 4713 (2021).

[24] C. Zheng, P. Thomas, and E. Tang, Robust topological oscillators govern a tunable phase transition to synchronized circadian rhythms, arXiv, doi:10.48550/arXiv.2607.13322 (2026), 2607.13322.

[25] M. Nakajima, H. Ito, and T. Kondo, In vitro regulation of circadian phosphorylation rhythm of cyanobacterial clock protein KaiC by KaiA and KaiB, FEBS Lett. 584, 898 (2010).

[26] J. Snijder, J. M. Schuller, A. Wiegard, P. Lössl, N. Schmelling, I. M. Axmann, J. M. Plitzko, F. Förster, and A. J. R. Heck, Structures of the cyanobacterial circadian oscillator frozen in a fully assembled state, Science 355, 1181 (2017).

[27] Y. Murayama, A. Mukaiyama, K. Imai, Y. Onoue, A. Tsunoda, A. Nohara, T. Ishida, Y. Maeda, K. Terauchi, T. Kondo, and S. Akiyama, Tracking and visualizing the circadian ticking of the cyanobacterial clock protein KaiC in solution, EMBO J. 30, 68 (2011).

[28] K. Oyama, C. Azai, K. Nakamura, S. Tanaka, and K. Terauchi, Conversion between two conformational states of KaiC is induced by ATP hydrolysis as a trigger for cyanobacterial circadian oscillation, Sci. Rep. 6, 32443 (2016).

[29] Y. G. Chang, N. W. Kuo, R. Tseng, and A. LiWang, Flexibility of the C-terminal, or CII, ring of KaiC governs the rhythm of the circadian clock of cyanobacteria, Proc. Natl. Acad. Sci. U S A 108, 14431 (2011).

[30] X. Han, D. Zhang, L. Hong, D. Yu, Z. Wu, T. Yang, M. Rust, Y. Tu, and Q. Ouyang, Determining subunit-subunit interaction from statistics of cryo-EM images: observation of nearestneighbor coupling in a circadian clock protein complex, Nat. Commun. 14, 5907 (2023).

[31] X. Qin, M. Byrne, T. Mori, P. Zou, D. R. Williams, H. McHaourab, and C. H. Johnson, Intermolecular associations determine the dynamics of the circadian KaiABC oscillator, Proc. Natl. Acad. Sci. U S A 107, 14805 (2010).

[32] Y. Kuramoto, Chemical Oscillations, Waves, and Turbulence (Dover, 1984).

[33] K. Kondo, Y. Furuike, K. Horiuchi, Y. Onoue, E. Yamashita, and S. Akiyama, Local and non-local impacts of intra-hexamer interactions on temperature compensation of KaiC, bioRxiv, doi:10.64898/2026.06.12.727564 (2026).

[34] H. Frauenfelder, G. Chen, J. Berendzen, P. W. Fenimore, H. Jansson, B. H. McMahon, I. R. Stroe, J. Swenson, and R. D. Young, A unified model of protein dynamics, Proc. Natl. Acad. Sci. U S A 106, 5129 (2009).

[35] V. Lubchenko, P. G. Wolynes, and H. Frauenfelder, Mosaic energy landscapes of liquids and the control of protein conformational dynamics by glass-forming solvents, J. Phys. Chem. B 109, 7488 (2005).

[36] V. Lubchenko, Quantitative theory of structural relaxation in supercooled liquids and folded proteins, J. Non-Cryst. Solids 352, 4400 (2006).

[37] D. Bell-Pedersen, V. M. Cassone, D. J. Earnest, S. S. Golden, P. E. Hardin, T. L. Thomas, and M. J. Zoran, Circadian rhythms from multiple oscillators: lessons from diverse organisms, Nat. Rev. Genetics 6, 544 (2005).

[38] P. Ruoff, J. J. Loros, and J. C. Dunlap, The relationship between FRQ-protein stability and temperature compensation in the Neurospora circadian clock, Proc. Natl. Acad. Sci. U S A 102, 17681 (2005).

[39] Y. Oono and M. Paniconi, Steady state thermodynamics, Progress of Theoretical Physics Supplement 130, 29 (1998).

[40] T. Hatano and S. Sasa, Steady-state thermodynamics of Langevin systems, Phys. Rev. Lett. 86, 3463 (2001).

[41] K. Yoshimura, A. Kolchinsky, A. Dechant, and S. Ito, Housekeeping and excess entropy production for general nonlinear dynamics, Phys. Rev. Res. 5, 013017 (2023).

[42] M. Esposito and C. Van den Broeck, Three detailed fluctuation theorems, Phys. Rev. Lett. 104, 090601 (2010).

[43] M. Esposito and C. Van den Broeck, Three faces of the second law. I. Master equation formulation, Phys. Rev. E 82, 011143 (2010).

[44] C. Van den Broeck and M. Esposito, Three faces of the second law. II. Fokker-Planck formulation, Phys. Rev. E 82, 011144 (2010).

[45] D. Zhang, Y. Cao, Q. Ouyang, and Y. Tu, Altruistic resource-sharing mechanism for synchronization: The energy-speed-accuracy trade-off, Phys. Rev. Lett. 135, 037401 (2025).

[46] H. Ito, H. Kageyama, M. Mutsuda, M. Nakajima, T. Oyama, and T. Kondo, Autonomous synchronization of the circadian KaiC phosphorylation rhythm, Nat. Struct. Mol. Biol. 14, 1084 (2007).

[47] M. J. Rust, S. S. Golden, and E. K. O’Shea, Light-driven changes in energy metabolism directly entrain the cyanobacterial circadian oscillator, Science 331, 220 (2011).

[48] T. Yoshida, Y. Murayama, H. Ito, H. Kageyama, and T. Kondo, Nonparametric entrainment of the in vitro circadian phosphorylation rhythm of cyanobacterial KaiC by temperature cycle, Proc. Natl. Acad. Sci. U S A 106, 1648 (2009).

[49] C. Zheng and E. Tang, A topological mechanism for robust and efficient global oscillations in biological networks, Nat. Comm. 15, 6453 (2024).

[50] H. Sakaguchi, Cooperative phenomena in coupled oscillator systems under external fields, Progr. Theor. Phys. 79, 39 (1988).

