## Supporting Text for "Molecular clockwork hypothesis for the KaiABC circadian oscillations" (from another version of this preprint: version 1, 2026-05-12)

### PACEMAKER OSCILLATIONS INDUCED BY ATPASE REACTIONS AND STRUCTURAL TRANSITIONS

We present a simple model to explain a potential mechanism by which the coupled dynamics of CI-ATPase reactions and structural transitions can induce oscillations. We define the probability of CI adopting the $p$ structure as $p^+$, and the probability of CI adopting the $dp$ structure as $p^- = 1 - p^+$. Additionally, we express the probability that CI binds ATP as $q^+ = \cos^2(\varepsilon/2)$, and the probability that CI binds ADP as $q^- = 1 - q^+ = \sin^2(\varepsilon/2)$. Therefore, $q^+ \approx 1$ when $\varepsilon \approx 2n\pi$, and $q^- \approx 1$ when $\varepsilon \approx (2n+1)\pi$, where $n$ is an integer.

We assume that the $p$ structure is stabilized when ATP binds to CI, while the $dp$ structure is stabilized when ADP binds to CI. In this framework, the release of Pi from CI is expected to trigger the transition from the $p$ structure to the $dp$ structure, which should occur at $\varepsilon \approx (2n + 1/2)\pi$ with $\sin\varepsilon \approx 1$. Similarly, the release of ADP, followed by the incorporation of ATP, should facilitate the transition from the $dp$ structure back to the $p$ structure, which is expected to happen at $\varepsilon \approx (2n + 3/2)\pi$ with $\sin\varepsilon \approx -1$. Using constants $a > 0$, $b > 0$, $c^+ > 0$, and $c^- > 0$, the stochastic process governing these structural transitions can be described by the following equation:

$$\frac{d}{dt}\begin{pmatrix} p^+ \\ p^- \end{pmatrix} = \begin{pmatrix} -a - c^+\sin\varepsilon & b - c^-\sin\varepsilon \\ a + c^+\sin\varepsilon & -b + c^-\sin\varepsilon \end{pmatrix}\begin{pmatrix} p^+ \\ p^- \end{pmatrix}. \tag{S1}$$

Additionally, we assume that the Pi release, which marks the transition from $q^+ \approx 1$ to $q^- \approx 1$, is promoted when CI adopts the $p$ structure with $p^+ \approx 1$, while the ADP release and the following ATP incorporation, which is the transition from $q^- \approx 1$ to $q^+ \approx 1$, is promoted when CI adopts the $dp$ structure with $p^- \approx 1$. Similar to modeling mechano-chemical coupling in molecular motors [1–3], we represent this scenario with the following Langevin equation:

$$\frac{d}{dt}\varepsilon = \omega_\varepsilon - (up^+ - vp^-)\sin\varepsilon + \xi_\varepsilon(t), \tag{S2}$$

where $u > 0$ and $v > 0$ are constants represnting the feedback effect of the CI structure on the reactions, and $\xi_\varepsilon$ represnts the noise effect, satisfying $\langle \xi_\varepsilon(t) \rangle = 0$ and $\langle \xi_\varepsilon(t)\xi_\varepsilon(t') \rangle = 2D_\varepsilon \delta(t - t')$, with a constant $D_\varepsilon > 0$. $\omega_\varepsilon > 0$ is the driving force of ATPase reactions with $\log(\omega_\varepsilon / D_\varepsilon) \propto (\mu_{\mathrm{ATP}} - (\mu_{\mathrm{ADP}} + \mu_{\mathrm{Pi}})) / (k_{\mathrm{B}} T)$, where $k_{\mathrm{B}}$ is the Boltzmann constant, $T$ is temperature, and $\mu_{\mathrm{ATP}}$, $\mu_{\mathrm{ADP}}$, and $\mu_{\mathrm{Pi}}$ are chemical potentials of ATP, ADP, and Pi, respectively. When $\omega_\varepsilon$ is sufficiently large, trapping of the reactions to $\varepsilon \approx n\pi$ is avoided, allowing cycles of reactions to be maintained. This presistence, in turn, drives the structural oscillations through the coupling shown in Eqs. S1 and S2. An example trajectory representing these oscillations is illustrated in Fig. S1. In this trajectory, the probability of ADP release (the probability to show $\sin \varepsilon \approx -1$) is large during the phosphorylation phase in the $p$ structure, which aligns with the experimental observations [4]. Further detailed simulations of the coupled dynamics of reactions and structural transitions in the KaiC hexamer were discussed in previous reports [5–8].

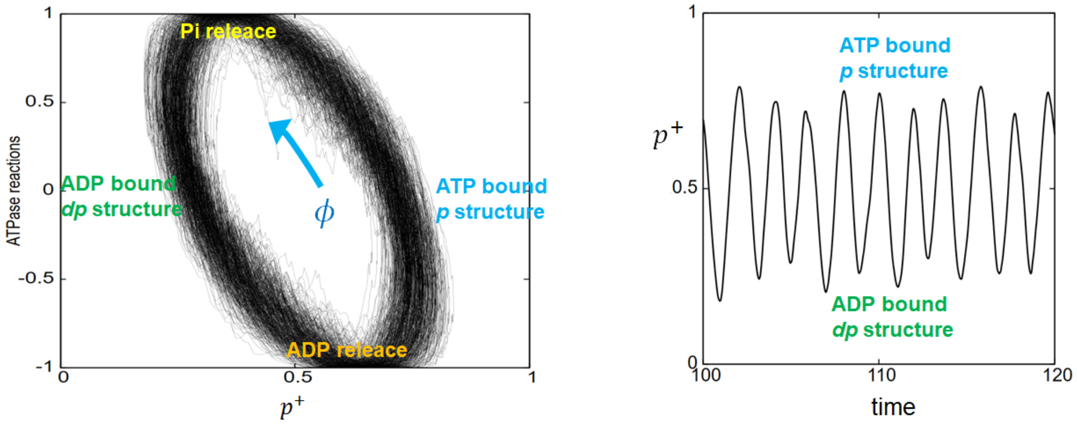


FIG. S1. Pacemaker oscillations induced by the coupled dynamics of ATPase reactions and structural transitions. (Left) An example trajectory of the simulated oscillations is plotted on the plane of the probability of exhibiting the $p$ structure, $p^+$, and the process occurring during the ATPase reactions, represented by $\sin \varepsilon$. The $p$ structure predominantly appeas when $p^+ > 0.5$, while the $dp$ structure becomes dominant when $p^+ < 0.5$. The phase of this limit cycle, $\phi$, iserves as the coordinate for the pacemaker oscillations discussed in the main text. (Right) The structural oscillations depicted in the left panel are shown as a function of time. Parameters used are: $a = b = c^+ = c^- = 1$, $u = v = 2$, $D_\varepsilon = 0.1$, and $\omega_\varepsilon = 3$.

**MEAN-FIELD FOKKER-PLANCK EQUATIONS**

From Eqs. 27 an 32 in the main text, the mean-field Fokker-Planck equations in the present study are

$$\frac{d}{dt}P_x(x) = -\frac{\partial}{\partial x}j_x(x)$$
$$j_x(x) = \left(\omega_x - b_x \sin x - D_x\frac{\partial}{\partial x}\right)P_x(x), \tag{S3}$$

and

$$\frac{d}{dt}P_y(y) = -\frac{\partial}{\partial y}j_y(y)$$
$$j_y(y) = \left(-\hat{\beta}_\phi r_\phi D_\phi \sin y - D_\phi\frac{\partial}{\partial y}\right)P_y(y), \tag{S4}$$

where $\hat{\beta}_\phi = \beta_\phi \langle\cos x\rangle / D_\phi$.

These two equations has the same form,

$$\frac{d}{dt}P_u(u) = -\frac{\partial}{\partial u}j(u)$$
$$j(u)/D = \left(a - b\sin u - \frac{\partial}{\partial u}\right)P_u(u). \tag{S5}$$

Eq. S3 for $u = x$ is recovered by putting $a = \omega_x/D_x$, $b = b_x/D_x$, and $D = D_x$, and Eq. S4 for $u = y$ is recovered by putting $a = 0$, $b = \hat{\beta}_\phi r_\phi$, and $D = D_\phi$. Eq. S5 has a static solution [9], satisfying $P_u(u) = P_u(u + 2\pi)$,

$$P_u(u) = \frac{h(u)}{Z}\left[1 + \left(e^{-2\pi a} - 1\right)\frac{g(u)}{g(2\pi)}\right], \tag{S6}$$

where

$$h(u) = \exp\left(ax + b\cos x\right), \qquad g(u) = \int_0^u dq \frac{1}{h(q)},$$
$$Z = \int_{-\pi}^{\pi} du h(u)\left[1 + \left(e^{-2\pi a} - 1\right)\frac{g(u)}{g(2\pi)}\right], \quad \text{and} \quad j_u = j_u^0 = -\frac{D}{Zg(2\pi)}\left(e^{-2\pi a} - 1\right). \tag{S7}$$

Using Eq. S6, expectation values are calculated as $\langle\cdots\rangle = \frac{1}{2\pi}\int_{-\pi}^{\pi} du(\cdots)P_u(u)$. Some of them can be expressed explicitly in the weak coupling ($b \ll 1$) and in the strong coupling ($b \gg 1$) cases as

- **Weak coupling case ($b \ll 1$)**

In this case, we have the following expressions:

$$\langle \cos u \rangle = \frac{1}{2(a^2+1)} b - \frac{1-2a^2}{4(a^2+1)^2(a^2+4)} b^3 + O(b^4), \tag{S8}$$

$$\langle \sin u \rangle = \frac{a}{2(a^2+1)} b - \frac{5a-a^3}{8(a^2+1)^2(a^2+4)} b^3 + O(b^4), \tag{S9}$$

$$\left\langle \cos^2 u \right\rangle = \frac{1}{2} + \frac{2-a^2}{8(a^2+1)(a^2+4)} b^2 + O(b^4), \tag{S10}$$

$$\left\langle \sin^2 u \right\rangle = \frac{1}{2} - \frac{2-a^2}{8(a^2+1)(a^2+4)} b^2 + O(b^4), \tag{S11}$$

$$\langle u \rangle = \frac{a}{a^2+1} b - \frac{3a}{4(a^2+1)(a^2+4)} b^2 + O(b^3), \tag{S12}$$

$$\sigma_u^2 = \left\langle u^2 \right\rangle - \langle u \rangle^2 = \frac{\pi^2}{3} - \frac{2}{a^2+1} b - \frac{5a^4+15a^2-2}{4(a^2+1)^2(a^2+4)} b^2 + O(b^3). \tag{S13}$$

- **Strong coupling case ($b \gg 1$)**

In this case, $P_u(u)$ can be approximated by a Gaussian,

$$P_u(u) \approx \frac{1}{\sqrt{2\pi\sigma_u^2}} \exp\left( -\frac{(u - \langle u \rangle)^2}{2\sigma_u^2} \right), \tag{S14}$$

where

$$\langle u \rangle = \arcsin\left(\frac{a}{b}\right), \tag{S15}$$

$$\sigma_u^2 = \left\langle u^2 \right\rangle - \langle u \rangle^2 = \frac{1}{b \cos \langle u \rangle} = \frac{1}{b\sqrt{1-\left(\frac{a}{b}\right)^2}}. \tag{S16}$$

Using this expression for $P_u(u)$, we have

$$\langle \cos u \rangle \approx e^{-\sigma_u^2/2} \cos \langle u \rangle = e^{-\sigma_u^2/2} \sqrt{1-\left(\frac{a}{b}\right)^2}, \tag{S17}$$

$$\langle \sin u \rangle \approx e^{-\sigma_u^2/2} \left(\frac{a}{b}\right), \tag{S18}$$

$$\left\langle \cos^2 u \right\rangle \approx \frac{1}{2}\left(1 + e^{-2\sigma_u^2} \cos \langle 2u \rangle\right) = \frac{1}{2}\left(1 + e^{-2\sigma_u^2}\left[1 - 2\left(\frac{a}{b}\right)^2\right]\right), \tag{S19}$$

$$\left\langle \sin^2 u \right\rangle \approx \frac{1}{2}\left(1 - e^{-2\sigma_u^2} \cos \langle 2u \rangle\right) = \frac{1}{2}\left(1 - e^{-2\sigma_u^2}\left[1 - 2\left(\frac{a}{b}\right)^2\right]\right). \tag{S20}$$

(S21)

**Transition from the non-oscillatory to oscillatory states**

The global ensemble-level oscillations are realized when $r_\phi \neq 0$. Here, the order parameter $r_\phi$ is defined by $\langle \cos \phi \rangle = r_\phi \cos \langle \phi \rangle$. Using $\phi = y + \omega t$, we have $\langle \cos(y + \omega t) \rangle =$

$\langle \cos y \rangle \cos \omega t - \langle \sin y \rangle \sin \omega t = r_\phi \cos \langle y \rangle \cos \omega t - r_\phi \sin \langle y \rangle \sin \omega t$. Because $\langle \sin y \rangle = 0$ and $\langle y \rangle = 0$, this definition becomes $r_\phi = \langle \cos y \rangle$. By putting $a = 0$ and $b = \hat{\beta}_\phi r_\phi$ into Eq. S8 for $u = y$, we have

$$r_\phi = \langle \cos y \rangle = \frac{1}{2}\hat{\beta}_\phi r_\phi - \frac{1}{16}(\hat{\beta}_\phi r_\phi)^3 + O\left((\hat{\beta}_\phi r_\phi)^4\right). \tag{S22}$$

This is the self-consistent relation to determine $r_\phi$. For $\hat{\beta}_\phi \leq 2$, $r_\phi = 0$ is the unique solution. When we neglect $O\left((\hat{\beta}_\phi r_\phi)^4\right)$ for $\hat{\beta}_\phi > 2$, we have another solution,

$$\begin{aligned} r_\phi &= \sqrt{\frac{8(\hat{\beta}_\phi - 2)}{\hat{\beta}_\phi^3}} \approx \sqrt{\hat{\beta}_\phi - 2} \\ &= \sqrt{\frac{\beta_\phi \langle \cos x \rangle}{D_\phi} - 2}. \end{aligned} \tag{S23}$$

Therefore, the increase in the coupling strength from $\hat{\beta}_\phi \leq 2$ to $\hat{\beta}_\phi > 2$ is the transition from the non-oscillatory to oscillatory states, where $r_\phi$ shows a critical behavior as in Eq. S23. For the larger coupling strength $\hat{\beta}_\phi \gg 1$, from Eq. S17, we find $r_\phi$ approaches to 1 as

$$r_\phi \sim \exp\left(-\frac{1}{2\hat{\beta}_\phi}\right) = \exp\left(-\frac{D_\phi}{2\beta_\phi \langle \cos x \rangle}\right). \tag{S24}$$

**Variances among oscillators**

When $r_\phi = 0$, we have $P_y(y) = 1/(2\pi)$, which corresponds to the $b = 0$ case in Eq. S13, resulting in $\sigma_\phi^2 = \pi^2/3$. When $r_\phi$ is positive, by putting $a = 0$ and $b = \hat{\beta}_\phi r_\phi$ in Eq. S13, we find $\sigma_\phi^2$ is reduced to $\sigma_\phi^2 \approx \pi^2/3 - 2\hat{\beta}_\phi r_\phi$. When the coupling $\hat{\beta}_\phi$ is large enough, from Eq. S16, we have $\sigma_\phi^2 \approx 1/b = D_\phi/(\beta_\phi r_\phi \langle \cos x \rangle)$.

$\sigma_\theta^2$ is calculated by

$$\sigma_\theta^2 = \langle (x+y) \rangle^2 - (\langle x + y \rangle)^2 = \sigma_x^2 + \sigma_\phi^2. \tag{S25}$$

When the couplings, $b_\theta$ and $b_\phi$, are large, we find $\sigma_x^2 \ll 1$ and $\sigma_\theta^2 \approx \sigma_\phi^2$.

**Entropy production rate at the steadily oscillating state**

In the mean-field expression, currents defined in Eq. 8 in the main text are

$$\begin{aligned} J_\theta &= \left(\omega_\theta - b_\theta \sin x - D_\theta \frac{\partial}{\partial x}\right) P_x(x) P_y(y), \\ J_\phi &= \left(\omega_\phi + b_\phi \sin x - \beta_\phi r_\phi \sin y \cos x - D_\phi \left(\frac{\partial}{\partial y} - \frac{\partial}{\partial x}\right)\right) P_x(x) P_y(y). \end{aligned} \tag{S26}$$

Using Eq. S26, the house-keeping [10] or the adiabatic [11–13] component of the entropy production rate at the steadily oscillating state is

$$\dot{S}_{\mathrm{a}} = \int d\theta d\phi \left( \frac{J_\theta^2}{P(\theta,\phi) D_\theta} + \frac{J_\phi^2}{P(\theta,\phi) D_\phi} \right), \tag{S27}$$

By putting $P(\theta,\phi) = P_x(x) P_y(y)$ into this relation, we have the following expression:

$$\begin{aligned} \dot{S}_{\mathrm{a}} =& \frac{\bar{\omega}^2}{\hat{D}} + 2\bar{\omega}\delta b \langle \sin x \rangle + \hat{D}\delta b^2 \left\langle \sin^2 x \right\rangle \\ &+ \frac{j_{0x}^2}{D_\theta + D_\phi} \int_{-\pi}^{\pi} \frac{dx}{P_x(x)} + \frac{\beta_\phi^2 r_\phi^2}{D_\phi} \left\langle \sin^2 y \right\rangle \left( \left\langle \cos^2 x \right\rangle - \langle \cos x \rangle^2 \right). \end{aligned} \tag{S28}$$

Here,

$$\begin{aligned} \hat{D} &= \frac{D_\theta D_\phi}{D_\theta + D_\phi}, \qquad \bar{\omega} = \hat{D} \left( \frac{\omega_\theta}{D_\theta} + \frac{\omega_\phi}{D_\phi} \right), \\ \delta b &= \frac{b_\phi}{D_\phi} - \frac{b_\theta}{D_\theta}, \qquad j_{0x} = -\frac{D_x}{Z_x g(2\pi)} \left( e^{-2\pi\omega_x/D_x} - 1 \right) \end{aligned} \tag{S29}$$

The term proportional to $\beta_\phi^2$ in Eq. S28 becomes zero when $r_\phi = 0$. This situation occurs either when the coupling $\beta_\phi$ is small, resulting in a loss of global synchronization, or when the couplings $b_\theta$ and $b_\phi$ are small, which leads to a small value of $\langle \cos x \rangle$. Conversely, when $\beta_\phi$, $b_\theta$, and $b_\phi$ are large, we anticipate $r_\phi \approx 1$ and $r_x = \langle \cos x \rangle / \cos \langle x \rangle \approx 1$. However, from Eqs. S17, S19, and S20, The term proportinal to $\beta_\phi^2$ becomes

$$\frac{\beta_\phi^2 r_\phi^2}{D_\phi} \left\langle \sin^2 y \right\rangle \left( \left\langle \cos^2 x \right\rangle - \langle \cos x \rangle^2 \right) \approx \frac{\beta_\phi^2 r_\phi^2}{D_\phi} \frac{\left(1 - r_\phi^4\right)\left(1 - r_x^2\right)}{2} \left[ 1 - r_x^2 \left( \cos^2 \langle x \rangle - \sin^2 \langle x \rangle \right) \right],$$

which also vanishes when $r_\phi \approx 1$ and $r_x \approx 1$. The other terms in Eq. S28 are calculated using the expressions for $P_x(x)$, $\langle \sin x \rangle$, and $\left\langle \sin^2 x \right\rangle$. To represent them explicitly, we can utilize Eqs. S9 and S11 in the weak coupling case, as well as Eqs. S18 and S20 in the strong coupling case.

In the weak coupling case, the term proportional to $j_{0x}^2$ in Eq. S28 becomes

$$\frac{j_{0x}^2}{D_\theta + D_\phi} \int_{-\pi}^{\pi} \frac{dx}{P_x(x)} \approx \frac{(\omega_\theta - \omega_\phi)^2}{D_\theta + D_\phi} \left( 1 - \frac{(b_\theta + b_\phi)^2}{(D_\theta + D_\phi)^2 A_0} \right), \tag{S30}$$

where

$$A_0 = 2 + 2 \left( \frac{\omega_\phi - \omega_\theta}{D_\theta + D_\phi} \right)^2.$$

Using Eqs. S9 and S11 in the first line of Eq. S28, retaining the terms up to $O(b_\theta^2/D_\theta^2)$ and $O(b_\phi^2/D_\phi^2)$, and combining it with the expression of Eq. S30, we have

$$\dot{S}_{\mathrm{a}} \approx \dot{S}_\omega + \dot{S}_b + \dot{S}_{\omega b},$$

where

$$\dot{S}_\omega = \frac{\omega_\phi^2}{D_\phi} + \frac{\omega_\theta^2}{D_\theta},$$
$$\dot{S}_b = \frac{\hat{D}}{2}\delta b^2,$$
$$\dot{S}_{\omega b} = \frac{(\omega_\phi - \omega_\theta)(b_\theta + b_\phi)}{D_\theta + D_\phi} G$$
$$G = -\frac{1}{A_0}\left[\frac{2D_\theta D_\phi \delta b}{(D_\theta + D_\phi)^2}\left(\frac{\omega_\phi}{D_\phi} + \frac{\omega_\theta}{D_\theta}\right) + \frac{(\omega_\phi - \omega_\theta)(b_\theta + b_\phi)}{(D_\theta + D_\phi)^2}\right], \tag{S31}$$

which leads to Eq. 14 in the main text.

In the strong coupling case, the term proportional to $j_{0x}^2$ in Eq. S28 vanishes because $j_{0x} \approx 0$. Then, using Eqs. S18 and S20 in the first line of Eq. S28, we have

$$\dot{S}_\mathrm{a} \approx \dot{S}_\omega + \dot{S}_b + \dot{S}_{\omega b} + \dot{S}_{\beta_\phi},$$

where

$$\dot{S}_\omega = x\hat{D}\left[\frac{\omega_\phi}{D_\phi} + \frac{\omega_\theta}{D_\theta} - \delta b\left(\frac{\omega_\phi - \omega_\theta}{b_\phi + b_\theta}\right)\right]^2, \tag{S32}$$
$$\dot{S}_b = 2(1 - r_x)\hat{D}\delta b^2, \tag{S33}$$
$$\dot{S}_{\omega b} = 2(1 - r_x)\hat{D}\delta b\left(\frac{\omega_\phi - \omega_\theta}{b_\phi + b_\theta}\right)\left[\frac{\omega_\phi}{D_\phi} + \frac{\omega_\theta}{D_\theta} - 2\delta b\frac{\omega_\phi - \omega_\theta}{b_\phi + b_\theta}\right], \tag{S34}$$
$$\dot{S}_{\beta_\phi} = 4(1 - r_\phi)(1 - r_x)\frac{\beta_\phi^2}{D_\phi}\left(\frac{\omega_\phi - \omega_\theta}{b_\phi + b_\theta}\right)^2. \tag{S35}$$

Then, in the strong coupling limit, neglecting the terms proportional to $1 - r_\phi$ or $1 - r_x$, we have $\dot{S}_\mathrm{a} \approx \dot{S}_\omega$. When the activity in the structural transitions driven by the CI-ATPase reactions is large with $\omega_\phi \gg \omega_\theta$, this expression leads to

$$\dot{S}_\mathrm{a} \approx \dot{S}_\omega \approx \frac{\omega^2}{\hat{D}}, \tag{S36}$$

which is Eq. 15 in the main text.

---


[1] R. D. Astumian and M. Bier, Fluctuation driven ratchets: Molecular motors, Phys. Rev. Lett. **72**, 1766 (1994).

[2] F. Jülicher, A. Ajdari, and J. Prost, Modeling molecular motors, Rev. Mod. Phys. **69**, 1269 (1997).

[3] D. Keller and C. Bustamante, The mechanochemistry of molecular motors, Biophys. J. **78**, 541 (2000).

[4] K. Terauchi, Y. Kitayama, T. Nishiwaki, K. Miwa, Y. Murayama, T. Oyama, and T. Kondo, ATPase activity of KaiC determines the basic timing for circadian clock of cyanobacteria, Proc. Natl. Acad. Sci. U S A **104**, 16377 (2007).

[5] J. Paijmans, D. K. Lubensky, and P. R. Ten Wolde, A thermodynamically consistent model of the post-translational Kai circadian clock, PLoS Comput. Biol. **13**, e1005415 (2017).

[6] S. Das, T. P. Terada, and M. Sasai, Role of ATP hydrolysis in cyanobacterial circadian oscillator, Sci. Rep. **7**, 17469 (2017).

[7] M. Sasai, Role of the reaction-structure coupling in temperature compensation of the KaiABC circadian rhythm, PLoS Comput. Biol. **18**, e1010494 (2022).

[8] M. Sasai, Mechanism of autonomous synchronization of the circadian KaiABC rhythm, Sci. Rep. **11**, 4713 (2021).

[9] H. Sakaguchi, Cooperative phenomena in coupled oscillator systems under external fields, Progr. Theor. Phys. **79**, 39 (1988).

[10] T. Hatano and S. Sasa, Steady-state thermodynamics of Langevin systems, Phys. Rev. Lett. **86**, 3463 (2001).

[11] M. Esposito and C. Van den Broeck, Three detailed fluctuation theorems, Phys. Rev. Lett. **104**, 090601 (2010).

[12] M. Esposito and C. Van den Broeck, Three faces of the second law. I. Master equation formulation, Phys. Rev. E **82**, 011143 (2010).

[13] C. Van den Broeck and M. Esposito, Three faces of the second law. II. Fokker-Planck formulation, Phys. Rev. E **82**, 011144 (2010).
